# Efflux-linked Accelerated Evolution of Antibiotic Resistance at a Population Edge

**DOI:** 10.1101/2022.04.01.486762

**Authors:** Souvik Bhattacharyya, Madhumita Bhattacharyya, Dylan M. Pfannenstiel, Anjan K. Nandi, YuneSahng Hwang, Khang Ho, Rasika M. Harshey

## Abstract

Efflux is a common mechanism of resistance to antibiotics. We show that efflux itself promotes accumulation of antibiotic-resistance mutations (ARM). This phenomenon was initially discovered in a bacterial swarm where the linked phenotypes of high efflux and high mutation frequencies spatially segregated to the swarm edge, driven there by motility. We uncovered and validated a global regulatory network that connects high efflux to downregulation of specific DNA repair pathways even in non-swarming states. The efflux-DNA repair link was corroborated in a clinical ‘resistome’ database: genomes with mutations that increase efflux exhibit a significant increase in ARMs. Accordingly, efflux inhibitors decreased evolvability to antibiotic resistance. Swarms also revealed how bacterial populations serve as reservoir of ARMs even in absence of antibiotic selection pressure. High efflux at the edge births mutants that, despite compromised fitness, survive there because of reduced competition. This finding is relevant to biofilms where efflux activity is high.

## INTRODUCTION

Antibiotic resistance is a global health concern and is being labeled a ‘silent pandemic’(Mahoney et al., 2021; Mendelson et al., 2022). Genetic antibiotic resistance can evolve by natural selection when bacteria are exposed to antibiotics (Shallcross et al., 2017; Traxler and Kolter, 2015). This exposure not only allows positive selection of pre-existing mutations but can also activate efflux pumps (Du Toit, 2017). These pumps are diverse, numerous, and critical for several bacterial physiologies including expulsion of intracellular antibiotics (Nishino et al., 2009). When challenged with antibiotic stress, bacteria activate efflux pathways almost ubiquitously (Webber and Piddock, 2003). ARMs (antibiotic-resistance mutations) that increase the efflux of antibiotics are widely described in clinical isolates (Nikaido and Pages, 2012). We report the existence of a genetic network that connects elevated efflux activity to the downregulation of DNA repair. This finding has clinical implications since antibiotic selection pressure in treatment regimens would induce efflux and consequently promote ARMs. Our analysis of a resistome database curated for ARMs in both clinical and non-clinical isolates showed a positive correlation between ARMs that increase efflux and total ARMs in the genome.

ARMs are vulnerable to loss in the absence of antibiotic selection pressure because the mutants invariably experience a reduction in fitness. This is because of the essentiality of the cellular targets of antibiotics - transcription, translation, or cell wall biogenesis – processes most often modified by the genetic resistance mechanisms (Kohanski et al., 2010). When competing with susceptible individuals in an antibiotic-free environment, mutations that confer resistance but alter cellular physiology are disadvantageous (Andersson and Levin, 1999; Bottger et al., 2005; Melnyk et al., 2015). This has been demonstrated to be especially true when the density of population (and hence competition) is high (Hibbing et al., 2010; Otto and Whitlock, 1997). And yet, they are seen to persist in dense populations such as biofilms (Steenackers et al., 2016). We report the discovery of a special ecology at the edge of a dense population where ARMs survive. Cells exhibiting high efflux and high mutation frequencies segregate to the edge, where they spawn such mutants.

Our model dense population is the *E. coli* swarm, where the bacteria move collectively over a semi-solid surface, powered by flagella (Harshey, 2003; Partridge and Harshey, 2013b). Cells in a swarm grow and divide as they colonize ever-increasing swaths of territory; these cells are physiologically distinct from their planktonic counterparts. Unlike dense biofilms, in which bacteria have reduced metabolism and slow growth (Flemming et al., 2016; Hoiby et al., 2010), swarms are metabolically highly active (Kearns, 2010; Partridge and Harshey, 2013a). An unexpected property of bacterial swarms is their tolerance to a broad class of antibiotics, which dissipates when the bacteria are removed from the swarm and dispersed in liquid (Butler et al., 2010; Kim et al., 2003; Lai et al., 2009). ‘Tolerance’ and ‘persistence’ are adaptive physiological mechanisms in bacteria that temporarily either turn down growth or turn up multidrug efflux pumps (Brauner et al., 2016). Swarms exhibit elevated efflux and ROS catabolism and decreased membrane permeability even in the absence of antibiotic exposure, suggesting perhaps an anticipatory program to survive an encounter with antibiotics. ROS stress is known to be generated by high cell densities (Chang et al., 2002; Dukan and Nystrom, 1998) and by increased energy utilization expected during efflux (Anes et al., 2015; Fang, 2011; Imlay, 2013). When confronted with antibiotics, swarms further elevate both efflux and ROS catabolism. One of the proximate causes of this elevation is release of a ‘necrosignal’ from antibiotic-induced death of a sub-population. The antibiotic tolerance-enhancing factor was identified as being a part of an efflux pump itself: AcrA, a periplasmic component of the AcrAB-TolC RND efflux pump, is released from dead cells and signals live cells by interacting with TolC in the outer membrane to enhance drug efflux as well as to induce expression of many other efflux pumps (Bhattacharyya et al., 2020). We have called this phenomenon ‘necrosignaling’ and have demonstrated its existence in other Gram-negative as well as Gram-positive bacteria (Bhattacharyya et al., 2020). High efflux in swarms has parallels with ARMs that increase efflux in clinical settings.

We report here the convergence of several phenomena that create a high-evolvability niche at the advancing edge of an *E. coli* swarm. Cells at the edge are engaged in high efflux, likely for bringing in hard-to-get iron on a surface terrain. High efflux generates ROS stress, which may play a role in activating a regulatory pathway that connects efflux to downregulation of specific DNA repair genes, producing a mutator phenotype. Motility plays another important role in assisting this phenotype to ‘surf’ to the edge of the swarm where chances of fixation of unfit mutations are expected to be high due to reduced competition. Edge cells incur more cell death as well, releasing the necrosignal AcrA, which stimulates efflux further. We show that the mutator phenotype can be suppressed by inhibitors that target both AcrA and the efflux pump.

## RESULTS AND DISCUSSION

### Swarms Downregulate DNA Repair and Show High Evolvability to Antibiotic Resistance

Compared to planktonic populations, *E. coli* swarms intrinsically upregulate efflux and ROS catabolism pathways (Bhattacharyya et al., 2020). A study showing a correlation between increased efflux pump activity and downregulation of the DNA repair gene *mutS* in planktonic cells of *E. coli* (El Meouche and Dunlop, 2018), led us at first to re-examine our *E. coli* RNA seq data from swarms, which showed a 3-6 fold downregulation of eight DNA repair genes – *dinB, holA, lexA, mug, mutM, mutS, recC*, and *ung* (Figure 1A inset and Figure S1A). These genes are broadly involved in mismatch and nucleotide-excision repair, predicting a mutator phenotype. This was tested by examining for the presence of mutants resistant to the antibiotic Kanamycin (Kan^R^) in the swarm, using a modification of the border-crossing assay originally designed to demonstrate antibiotic tolerance in swarms (Butler et al., 2010). *E. coli* can swarm collectively on the surface of media solidified with ~0.5% agar (videos S1, S2), but will swim as individuals within softer agar (0.3%) (Partridge and Harshey, 2013a). Swimming bacteria are effectively in a planktonic state, so we will henceforth use S and P symbols to denote swarm and planktonic states. In the assay, a 2-chamber Petri dish filled with swarm media contains antibiotic in only one chamber; bacteria that begin swarming in the no-antibiotic chamber and arrive at the separating border, will attempt to cross over and colonize the antibiotic chamber. To detect individual bacteria harboring Kan^R^ mutations that arose within the swarm, we modified this assay to contain swim media with Kan in the right chamber (Figure 1B schematic; S to P setup). A control set-up for observing such mutants arising in the P population had swim media in both chambers (P to P setup).

**Figure 1.**
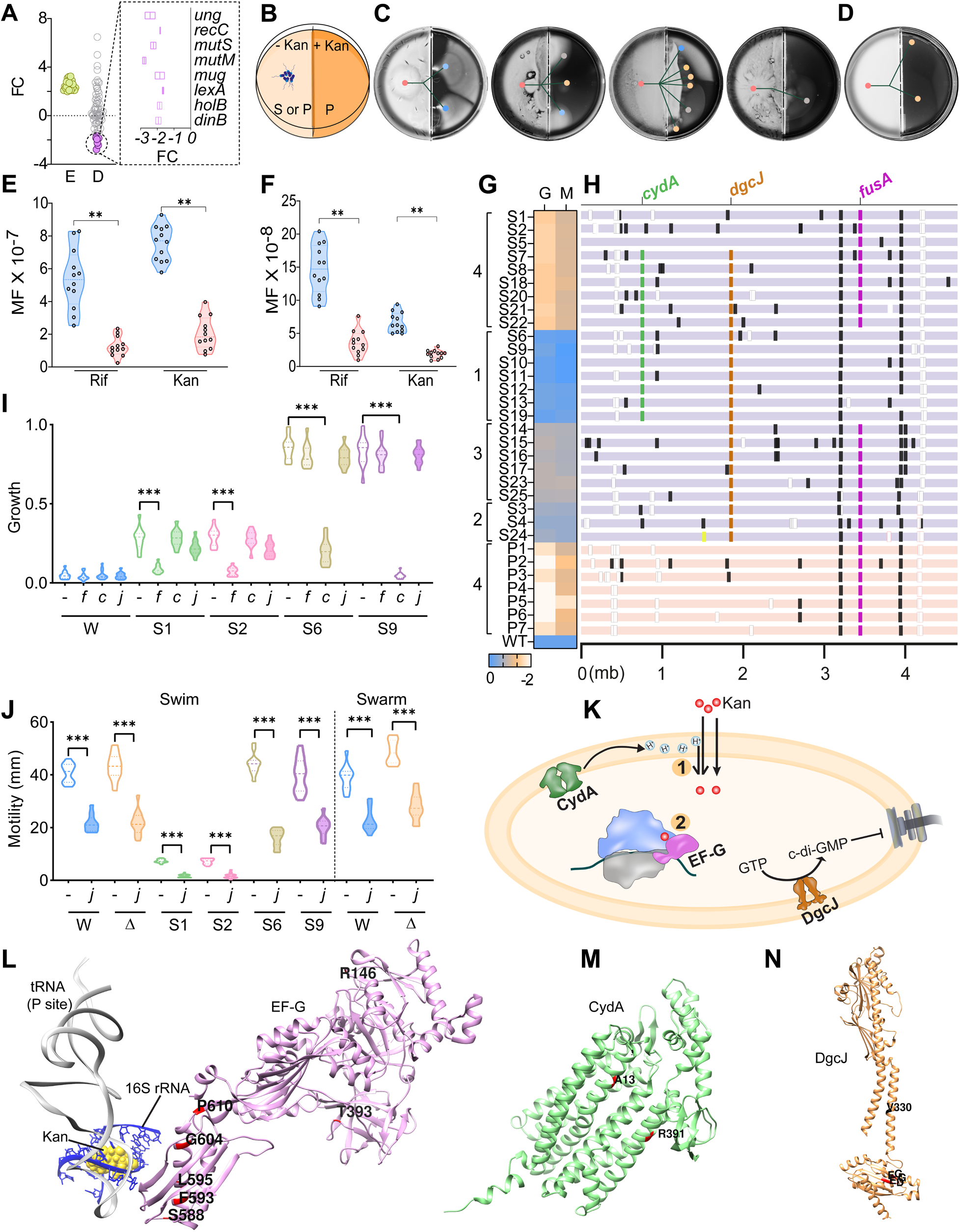
Swarms Downregulate DNA Repair and Show High Evolvability to Antibiotic Resistance. (A) Log2-fold change (FC) in expression of efflux and DNA-repair genes in *E. coli* swarm cells; E: efflux, D: DNA repair. Inset: DNA-repair genes downregulated more than 1.5 log2-fold. (B) Border-crossing assay schematic. S: swarm, P: swim; Kan = 25u/ml Kanamycin or Kan^25^. Bacteria are inoculated in the left no-Kan chamber. (C-D) WT *E. coli* inoculated on the left on either swarm (C) or swim (D) media, crossed the border into Kan^25^ swim media. Representative plates show the emergence of Kan-resistant (Kan^R^) swimmers. Color-coded circles mark distinct phenotypes (see H): Lines between circles indicate the lineage of mutants from WT. (E-F) Mutation frequencies (MF) of S (blue) and P (red) cells of (E) *E. coli* and (F) *B. subtilis*, measured by resistance to Rifampicin (Rif^50^) and Kan^50^. Each black circle is an experimental replicate. [\*\**p* < 0.01, Mann-Whitney test] (G) Heat map showing the growth G, and motility M, of Kan^R^ mutants from C-D, compared to WT. S1-25 were derived from C and P1-7 from D. The clusters of isolated mutants are indicated as 1-4 matching mutant color codes from (C-D)). (H) Summary of mutations from the individual isolates listed in G scaled to the *E. coli* genome in Mb (bottom). Syntenic mutations in the 3 indicated genes were recovered frequently. White and black boxes = inter- or intragenic mutations. (I-J) Comparison of growth (I) of indicated mutant and WT strains in Kan^25^ media, and motility (J) in swim media. *f, c, g* indicates complementation with plasmids expressing *fusA, cydA, dgcJ;* Δ, Δ*dgcJ*. [\*\*\**p* < 0.001, Wilcoxon rank sum test] (K) A schematic summarizing possible effects of recovered mutations in *fusA*, *cydA*, *dgcJ*. (1) *cydA* mutations reduce PMF which may decrease Kan uptake. (2) *fusA* mutations reverse the translation inhibition by Kan. *dgcJ* mutations are expected to reduce c-di-GMP and increase motility. (L-N) Structures of (L) EF-G (*fusA*; magenta) [PDB:4V7D], (M) CydA [PDB:6RKO], and (N) DgcJ [Alphafold]. Red: mutations in active site (residues 423-427); black: mutations in the predicted sensory domain. In each gene, multiple mutations from different S-P isolates map to the same site.

Kan^R^ mutants were recovered in both P-P and S-P set-ups (Figure 1C-D). The distinct points along the border from where the Kan^R^ swim ‘flares’ emerge report on the physical location of the original mutant in the S or P population, which then colonizes the area on the right. Kan^R^ mutants were recovered at a higher frequency per plate in S-to-P compared to P-to-P setups (Figure 1C-D and Figure S1B). A total of 32 Kan^R^ mutants were collected, 25 that arose from S-P setup, and 7 from the P-P setup. To determine the mutation frequency (MF) accurately, we measured resistance to the antibiotics Rifampicin (Rif) and Kan (Fowler et al., 1994; Kapoor et al., 2019; Miller et al., 1977) in both populations. Compared to P cells, S cells of *E. coli* exhibited a ~5-fold higher MF (Figure 1E). To test if this was a general feature of S cells, MFs were also calculated for S cells of *Bacillus subtilis* (Figure 1F) and *Pseudomonas aeruginosa* (Figure S1C), which showed ~8- and ~7-fold MF increases, respectively, compared to their P counterparts.

Differences in the intensity and size of the flares that emerged in the *E. coli* set up in Figure 1C, could be indicative of differences in the proficiency of growth and/or motility of the mutants. We therefore examined these parameters closely in all 32 mutants, as well as identified the mutations by whole-genome sequencing. Mutants derived from P cells (P1-7) were uniformly poor in motility (M) (Figure 1G), whereas those from S cells varied in this property, some moving even better than wild type (WT) (S1-25) (Figure 1G and Figure S1D). P-derived mutants were also uniformly poor in growth (G) (group 4 in Figure 1G and Figure S1E), whereas S-derived mutants clustered into four distinct groups, some growing as well as WT (groups 1-4 in Figure 1G and Figure S1E). DNA sequencing revealed an enrichment of mutations in three genes – *fusA, cydA, dgcJ* - across the mutant set (Figure 1H). *fusA* encodes elongation factor G (EF-G), which is a known source of Kan^R^ mutants (Voorhees and Ramakrishnan, 2013) as Kan inhibits translation by binding to the A site in the 30S ribosomal subunit, blocking EF-G catalyzed translocation. *cydA* encodes subunit I of cytochrome bd-I terminal oxidase. Kan^R^ mutations previously isolated in *cydA* (Hol et al., 2016) resulted in decreased PMF and were inferred to have decreased Kan uptake (Becker and Cooper, 2013). The third gene *dgcJ* is a putative cyclic di-GMP synthase and has no reported connection to Kan^R^, but is expected to increase motility (Jenal et al., 2017; Pesavento et al., 2008). To test if these mutations directly conferred the Kan^R^ phenotype, wild-type copies of all three genes were overexpressed in four representative isolates, yielding the following results. Overexpression of *fusA* caused Kan^R^ to diminish only in strains with *fusA* mutations (Figure 1I, S1&S2 set; Figure S1F), while *cydA* overexpression reduced Kan^R^ only in strains with *cydA* mutations (Figure 1I, S6&S9 set; Figure S1F). Overexpression of *dgcJ* did not abrogate the Kan^R^ phenotype in strains with *dgcJ* mutations (Figure 1I, S6&S9 set), but greatly reduced swimming motility of all strains tested, including that of WT (Figure 1J, swim; Figure S1G), consistent with the known role of c-di-GMP (the enzymatic output of DgcJ) in inhibiting motility. In summary, these experiments confirm that mutations in *fusA* and *cydA* confer Kan^R^, whereas DgcJ may have a specific role in swarming motility as indicated by the abundance of *dgcJ* mutants in isolates only from the S samples.

The relationship of *cydA*, *fusA* and *dgcJ* mutations in promoting Kan^R^ and motility is sketched Figure 1K, based on literature reports. Mapping the mutations associated with the three genes onto the known or predicted structures of these three proteins showed that for EF-G, most mutations mapped to residues known to affect the translocation of the ribosome (Zhou et al., 2014), but the two that map to domain I and II of EF-G, respectively (Connell et al., 2007), have no known connection with translocation (R146 and T393) (Figure 1L). For CydA, the two mutants isolated were at positions not previously associated with Kan^R^ (Allison et al., 2011) (Figure 1M). For DgcJ, four were in the predicted enzyme active site (Hengge, 2009), while one mapped to the predicted periplasmic sensory domain (Figure 1N).

In summary, the results in this section show that *E. coli* swarms downregulate a set of eight specific DNA repair genes and expectedly, show higher MFs. This observation is not unique to *E. coli* swarms, as high MFs are also observed in *B. subtilis* and *P. aeruginosa* swarms. A large majority of Kan^R^ mutants recovered from edge of *E. coli* swarms are accompanied by mutations that improve their motility in the swarm.

### Downregulation of Mismatch and Base Excision Repair Pathways Contributes to the Higher MFs of Swarm Cells

A detailed analysis of all the mutations recovered in the genomes of the 32 mutants (Figure 1G) revealed that of 501 total mutations identified, A→G and T→C changes were each ~5% lower in S compared to P isolates, whereas T→G and C→T changes were each ~2-3% higher in S (Figure 2A and Figure S2A). Overall, transition mutations decreased by ~11% and transversions increased by ~8% in S isolates (Figure 2B, whole genome). Considering only the mutations in genes conferring Kan^R^ – *fusA* and *cydA* – transitions decreased ~ 21% and transversions increased ~6% in S compared to P cells (Figure 2B, Kan^R^). Two other types of mutations isolated across the genome were indels. Of these, insertions ranging from 4 −11 bp were found in *dgcJ* whereas deletions from 6-9 bp were found in *cydA*. We found no bias in the distribution of mutations in coding or non-coding regions (Figure 2B and Figure S2B).

**Figure 2.**
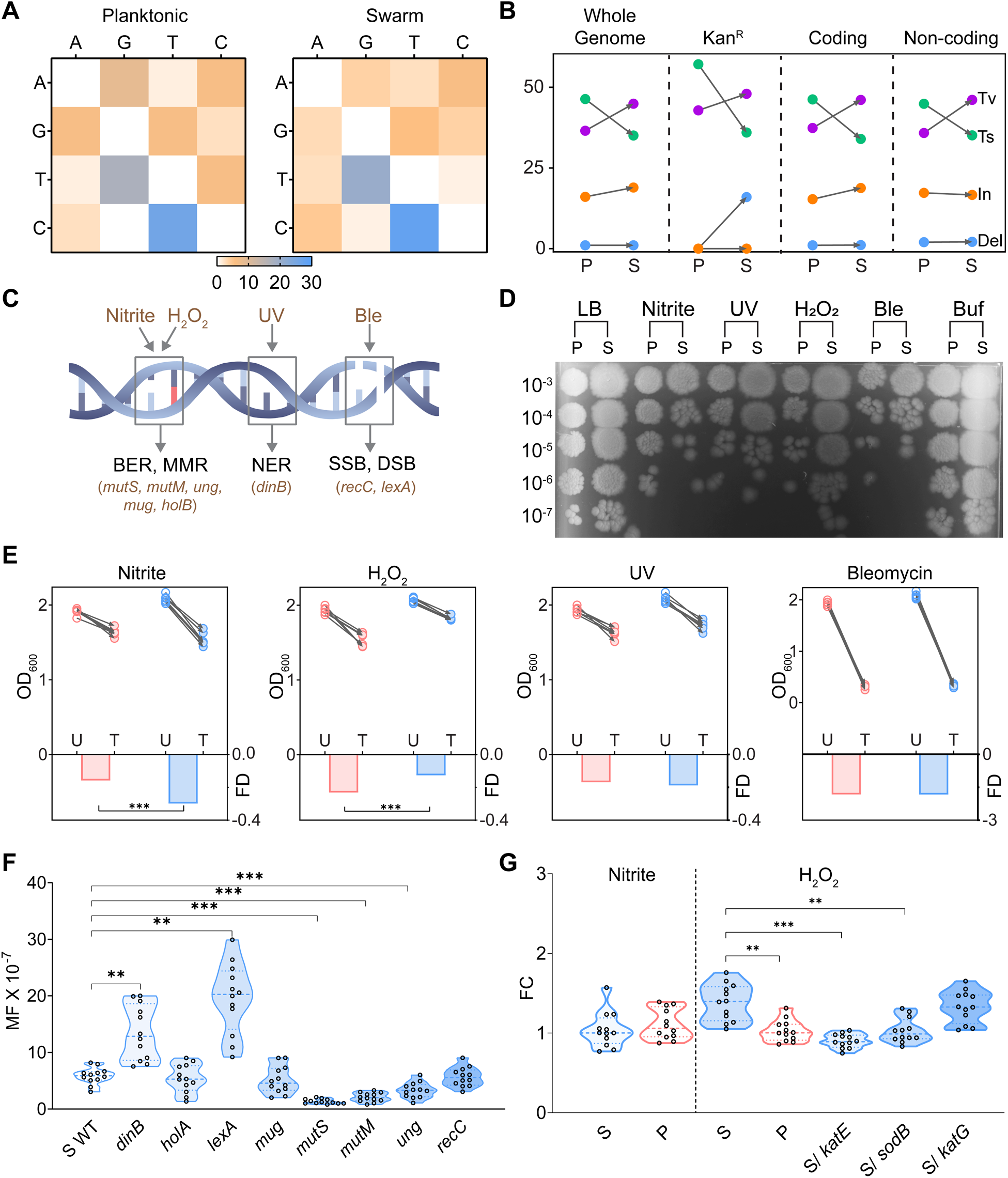
Downregulation of Mismatch and Base Excision Repair Pathways Contributes to the Higher MFs of Swarm Cells. (A) Heatmaps showing percentages of different types of mutations found in P and S mutants. WT nucleotides are on the left column, and rows show mutations in P1-P7 and S1-S25 isolates (Figure 1G). (B) A comparison of the percent changes (Y-axis) in types of mutations in S and P isolates. Transitions (Ts), Transversions (Tv), Insertions (In) and Deletions (Del) are color coded as indicated. All identified mutations in each group were first combined together in a set, then sorted on the basis of categories indicated on the top. (C) Schematic showing specific DNA repair pathways and genes that primarily function in these pathways. The major types of DNA damage caused by mutagenic agents used in this study are shown on top and the pathways (and genes) for dealing with them shown at the bottom of the DNA schematic. (D) Dilution spotting of S and P cells after treatment with: sodium nitrite (Nitrite), ultraviolet light (UV), hydrogen peroxide (H2O2), and bleomycin (Ble). (E) Comparison of cell growth (OD600) in the presence (T, treated) or absence (U, untreated) of indicated stresses. Blue: S, red: P. FD: mean fold-decrease in growth in T calculated as FD= log(T/U). [\*\**p* < 0.01, paired t-tests]. (F) Rif^50^ MF of S cells measured as in Figure 1E. WT S cells were transformed with clones of the indicated DNA repair genes and changes in MF monitored. \*\**p* < 0.01, \*\*\**p* < 0.001; Mann-Whitney test. (G) Fold changes (FC) in Nitrite and Peroxide levels in P and S cells, measured using commercial kits. Only peroxide levels were different. *katE, sodB* and *katG* genes were overexpressed in S cells to monitor changes in these levels. \*\**p* < 0.01, \*\*\**p* < 0.001; Wilcoxon rank-sum test.

To test which repair pathway might be responsible for the mutations recovered in S cells, these cells were exposed to mutagenic agents handled by the various pathways (Figure 2C). Acidified nitrite, a stressor for base excision, nucleotide excision, and mismatch repair pathways (Davies et al., 2011; Kurthkoti et al., 2008), significantly reduced the growth of S cells (Figure 2D-E). On the other hand, S cells survived better under the stress of hydrogen peroxide, an ROS agent affecting multiple DNA repair pathways (Figure 2D-E), consistent with the elevated expression in S cells of genes involved in scavenging ROS (Bhattacharyya et al., 2020). Bleomycin and UV light exerted significant but similar growth reduction in both P and S cells (Figure 2D-E) suggesting that DNA break repair pathways are not likely to be altered in S cells. We infer from these results that the downregulation of mismatch repair (MMR) and base excision repair (BER) pathways are likely contributing to the higher MF of S cells.

To further identify the specific pathways whose malfunction may have produced these mutations, we focused on the eight DNA repair genes (Friedberg, 2006; Rosenberg et al., 2012) downregulated in S cells (see Figure 1A), and overexpressed them using clones selected from the *E. coli* ASKA ORF-eome. Only clones of *mutS, mutM*, and *ung* significantly reduced MFs, which were now comparable to the MF of P cells (Figure 1E and 2F); MutS is an MMR protein, MutM is a formamidopyrimidine-DNA glycosylase and Ung is an uracil-DNA glycosylase. The latter two are involved in BER pathways (Figure 2C).

To identify the source of the damage contributing to the high MF of S cells, we monitored intracellular nitrite and peroxide levels using commercial kits. Levels of nitrite were similar between S and P cells, while intracellular peroxide levels were elevated in S cells (Figure 2G). Introduction of clones of known ROS catabolism genes *katE* and *sodB*, lowered the peroxide levels in S cells (Figure 2G). These results imply that despite the increase in expression of ROS catabolism genes observed in S cells (Bhattacharyya et al., 2020), these cells still experience ROS stress.

In summary, three lines of evidence – the nature of mutations recovered in S cells, the susceptibility of S cells to mutagens handled by MMR and BER pathways, and the elevated cellular peroxide levels in these cells, suggests that the high MF of S cells likely arises from both high cellular redox stress and decreased expression of the MMR and BER genes *mutS, mutM* and *ung*, which are known to handle DNA damage by ROS.

### A Global Regulatory Connection Between Efflux and DNA Repair

Efflux pump expression is tightly regulated by a vast network of regulators (Sun et al., 2014). To determine if efflux and DNA repair networks (Milanowska et al., 2011) were genetically connected, we took the bottoms-up approach summarized in Figure 3A. The regulatory connections between genes (nodes) in these two sets of pathways were interrogated by performing a nodal analysis of the three genes uncovered in this study - *ung*, *mutS*, and *mutM* - taken from the *E. coli* MG1655 curated genetic network Ecocyc (Keseler et al., 2011). We searched for regulatory paths connecting these three peripheral nodes to the efflux pump gene network, validating them by correlation of expression of gene-pairs using data from COLOMBOS [normalized *E. coli* gene expression data derived from 4000 experimental sets; (Moretto et al., 2016)]; we used our RNA seq data (Bhattacharyya et al., 2020) as an additional guide (Figure 3B-C and Figure S3A). From all the genes participating in these regulatory paths (Figure 3D top), only those gene pairs showing significant correlation coefficients from this analysis were considered (Figure 3C top), and are highlighted in Figure 3D (bottom) (see Figure S3B for paths that failed validation). The analysis revealed that downregulation of *mutS* and *ung* was controlled by *rpoS* via *sdsR* and *cpxR* nodes respectively, while downregulation of *mutM* was controlled by *crp* via *rpoH*. Therefore, *rpoS* and *crp* were the major nodes seen to connect efflux and DNA repair. RpoS is an alternative sigma factor, which controls the expression of hundreds of genes that protect the cell from stress and help survive nutrient deprivation (Landini et al., 2014), while CRP is a global transcription regulator of many genes mostly involved in energy metabolism (Grainger et al., 2005). We note that our analysis used datasets from thousands of experimental conditions not specific to *E. coli* swarms, so the identified links are evolutionarily wired and the consequences of their perturbation should be relevant in most niches these bacteria occupy.

**Figure 3.**
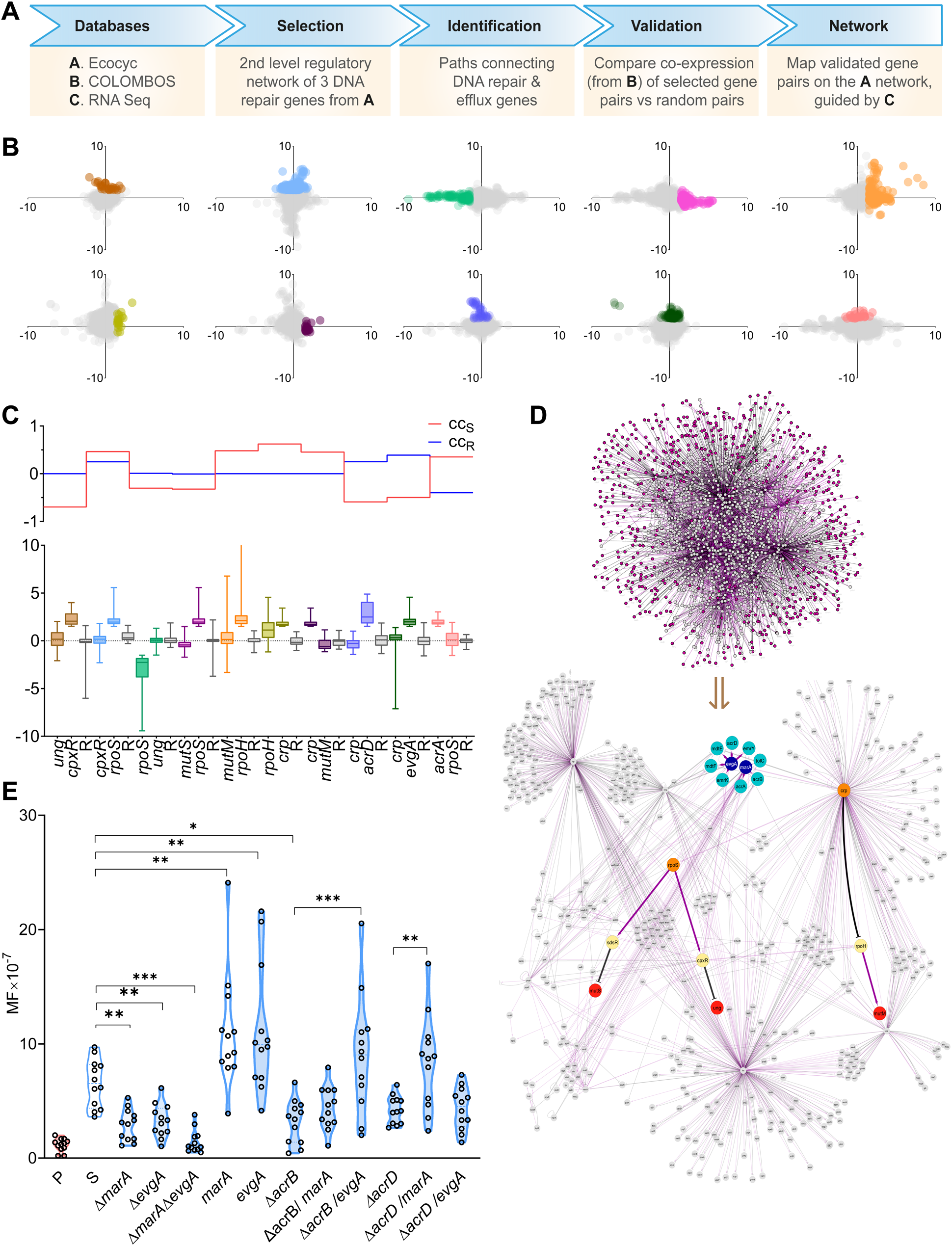
A Global Regulatory Connection Between Efflux and DNA Repair. (A) Flowchart summarizing the network analysis performed in this study. (B) Scatterplots showing expression of different gene-pairs within the network that connect efflux genes with DNA repair genes (color-coded as the gene-pairs in (C)); significant fold changes (± 1.5) are shown in matching colors with (C). (C) Bottom: Boxplot showing distribution of significant changes color-coded in B; R is one of 100 randomized pairs (see Figure S3A). Top: Lines indicating the calculated Pearson correlation coefficients (CC) of the corresponding pairs at the bottom. CC_S_, selected pair; CC_R_, random pair. (D) Top: Connection of 2^nd^ level regulatory network of DNA repair genes *mutS, mutM* and *ung* to efflux genes, selected from the 1^st^ level whole network (from database A in Figure 3A). Purple: terminal nodes; Grey: connecting nodes. Bottom: Validated portion of network from Top, with genetic connections between efflux (cyan) and DNA repair (red) genes. The connectors are efflux regulators (navy), major regulators of repair genes (orange), and genes connecting regulators with DNA repair (yellow). Dashed blue lines indicate an effect of efflux activity on regulators. Flatheads, downregulation; Arrows, upregulation. (E) Rif^50^ MF analysis of P (red) and S (blue) cells. X axis lists genetic background of S cells. [\**p* < 0.05, \*\**p* < 0.01, \*\*\**p* < 0.001; Mann-Whitney test]

What upstream signal might be relayed via the *rpoS* and *crp* nodes? High efflux pump activity is associated with various stress responses (Rosner and Martin, 2013). Efflux consumes an enormous amount of energy [ATP, PMF or Na+/H+ antiport systems drive various pumps (Du et al., 2018)] (Fang, 2011; Imlay, 2013). Like efflux, flagella biosynthesis and rotation during swarming also consume both PMF and ATP (Chevance and Hughes, 2008; Gabel and Berg, 2003). A high metabolic rate would be required to sustain a high PMF, which in turn would lead to high intracellular ROS. This was already inferred from gene expression data (Bhattacharyya et al., 2020), and demonstrated experimentally as well (Figure 2G). ROS are also produced by high cell densities (Chang et al., 2002; Dukan and Nystrom, 1998). Could the two global regulators RpoS and Crp identified in our analysis likely respond to ROS stress? This hypothesis was tested by measuring the effect on MF of deleting some of the regulators identified in the nodal analysis (*rpoS* and *crp* themselves were not deleted to avoid pleotropic effects (Ishihama et al., 2016)). Deletion of the major efflux regulators *marA* and *evgA* (Du et al., 2018) reduced MF, while their overexpression increased it (Figure 3E). A double deletion of *marA* and *evgA* decreased MF of S cells almost to the level of P cells (Figure 3E). The MarA regulon increases the expression of many efflux genes including *acrA, acrB*, and *tolC*. Similarly, EvgA controls the expression of efflux genes *acrD* and *mdtEF* (Du et al., 2018). To test whether the effect of these regulators on MF are pleiotropic or whether they occur via the genes they are known to regulate, we overexpressed *marA* in an Δ*acrB* strain, and *evgA* in an Δ*acrD* strain; in neither case did MF increase, showing that MarA and EvgA effects are routed through their specific genetic circuits. We conclude from both the nodal analysis and the experimental data presented, that the increased activity of efflux pumps itself downregulates specific DNA repair genes.

In summary, the experiments in this section show that the global regulators RpoS and Crp link specific DNA repair genes to efflux pump regulators. The results imply that these genetic links have been integrated into a network through evolutionary forces that increase mutagenesis and hence evolvability when bacteria experience high efflux-related ROS stress – an evolutionary model similar to other widely reported modes of stress-induced mutagenesis (Fitzgerald et al., 2017; MacLean et al., 2013; Ram and Hadany, 2014). Given that high efflux is a temporary state operative only during swarming, we suggest that the ultimate cause (Laland et al., 2011; Mayr, 1961), i.e. long-term or evolutionary importance of this reversible state is acquisition of genetic resistance.

### The Efflux-DNA Repair Network is Clinically Relevant to the Rise in Antibiotic Resistance

We have shown thus far that high-efflux leads to a mutator phenotype, which in turn creates genetic resistance to antibiotics or ARMs. Do these findings have any clinical significance? Mutations that increase efflux are known to provide resistance to antibiotics (Li et al., 2015). However, the MF data for clinical isolates are scarce. Based on the findings in Figure 3, we can expect to find a linkage between mutations that increase efflux and accumulation of ARMs in genomes of bacteria. To examine this, we used the Comprehensive Antibiotic Resistance Database (CARD) (Alcock et al., 2020), which catalogues ARMs for bacteria from both clinical and non-clinical sources. We analyzed only the resistome of *E. coli* (Figure 4A, Figure S4A-B). The data showed that the occurrence of efflux ARMs per genome follows a distribution pattern that is completely different from that of non-efflux ARMs. The fraction of genomes with efflux ARMs peaked at 17 mutations per genome and showed a gradual decrease up to 42 mutations per genome, but the fraction with non-efflux ARMs peaked at 1 mutation and started a steep decline at 4 mutations per genome (Figure 4B). More importantly, a strong positive correlation was found between efflux ARMs vs total ARMs in a genome (Figure 4C), supporting our hypothesis that efflux-promoting mutations tend to lead to more ARMs. Similar calculations with other categories of mutations did not show this association (Figure 4D-F, Figure S4C-E). A Spearman correlation matrix among these categories shows efflux and total mutations to be the only significantly correlated categories (Figure 4G, Figure S4F). These data suggest that, like in case of S cells, high efflux increases the rate of evolution of ARMs even in clinical settings, and consequently multidrug resistance.

**Figure 4.**
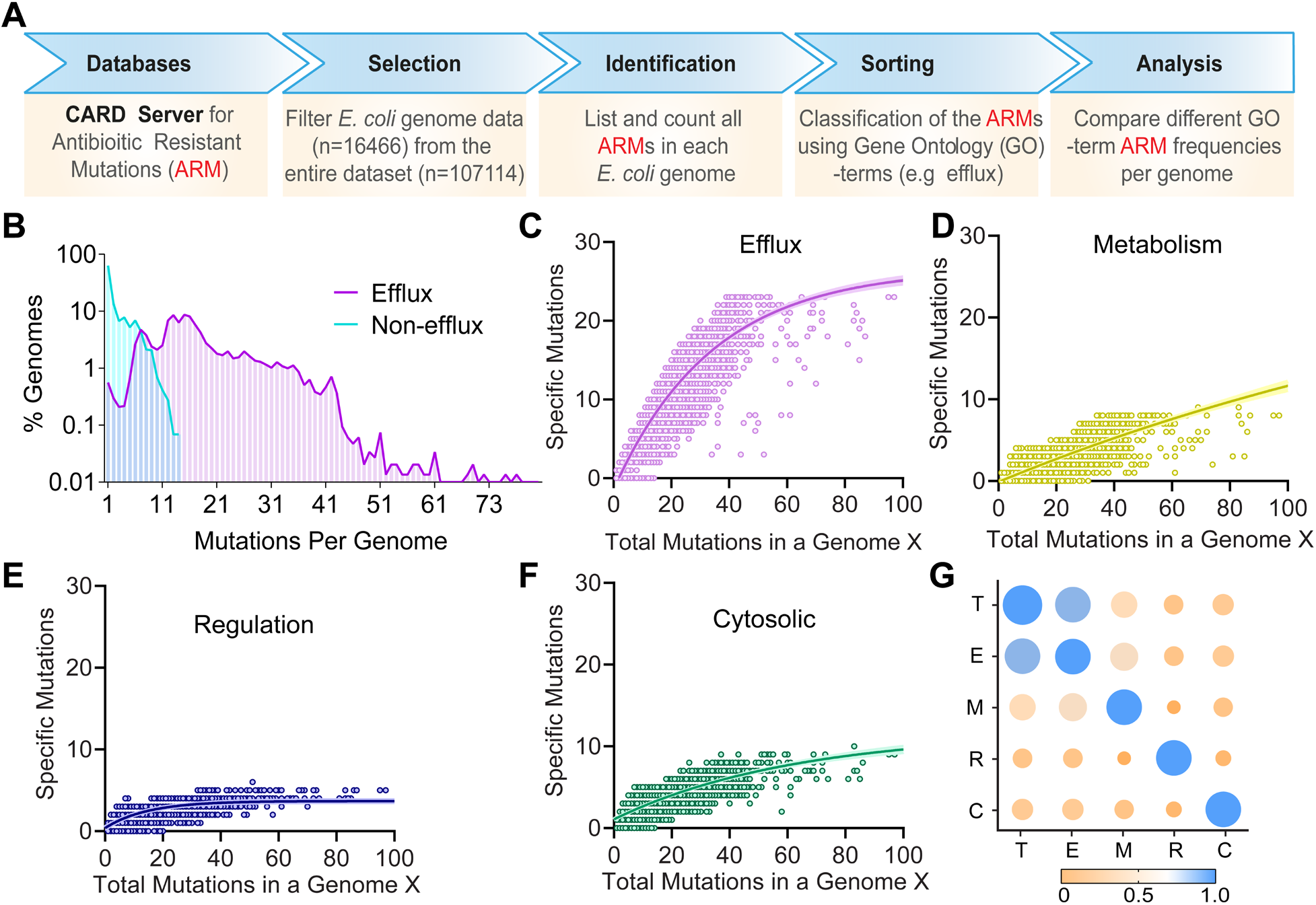
The Efflux-DNA Repair Network is Clinically Relevant to the Rise in Antibiotic Resistance. (A) Flowchart summarizing the analysis of the *E. coli* antibiotic resistome data from CARD. (B-F) Results from the analysis in (A). (B) Frequency distributions of mutations per genome in efflux and non-efflux genomes. (C-F) Scatter plots of total number of mutations in a given genome X vs number of ARMs in a specific category of genes (indicated on top) in that genome. One category from each parent GO (Gene Ontology) class was taken for comparison (see Figure S4A-B for details). (G) Bubble matrix of calculated Spearman correlation coefficients using the data in Figure 4C-F. T: total; E: Efflux; M: metabolism; R: regulatory; and C: cytosolic.

### The Mutator Phenotype of Swarms Can be Inhibited by Efflux Pump Inhibitors

The high efflux of swarm cells is further stimulated by the necrosignal AcrA, released upon death of a subpopulation (Bhattacharyya et al., 2020). It follows that AcrA addition should increase MF. We therefore incubated both P and S cells with purified AcrA and observed a significant increase of MF in S cells, but only a modest increase in P cells (Figure 5A). Conversely, inhibiting AcrA, which is an essential periplasmic component of the AcrA-TolC RND efflux pump, should not only reduce antibiotic outflow, but should also suppress the mutator phenotype. Using as a guide a recent study which identified several small molecules that target AcrA (Abdali et al., 2017), we selected three compounds – Novobiocin (Nov), Clorobiocin (Clo), and NSC60339 (Figure 5B-C) to study their effect on MF. Nov and Clo are both aminocoumarins that also target DNA gyrase activity (Gross et al., 2003) To circumvent this effect, we overproduced in the cells a *gyrB* mutant (R136L) conferring resistance to both Nov and Clo (Gross et al., 2003) (Figure S5A-C). We then treated both WT and Δ*acrA* S cells expressing *gyrB*^R136L^ with all three inhibitors in the presence or absence of externally added AcrA. Inhibition of AcrA was observed to decrease MF for all three compounds tested (Figure 5D-E). The increase of MF seen in WT S cells (i.e. without the *gyrB* allele) treated with Nov and Clo alone (Figure 5D, WT columns) is consistent with the idea that antibiotics increase DNA damage from redox stress (Dwyer et al., 2014) (Imlay, 2013). In contrast to Nov and Clo, the compound NSC60339 that solely targets AcrA to inhibit both efflux and necrosignaling, decreased MF even in WT. Among the three compounds tested, NSC60339 showed the best activity in decreasing both efflux- and necrosignal-mediated MF in all strain backgrounds (Figure 5D, NSC).

**Figure 5.**
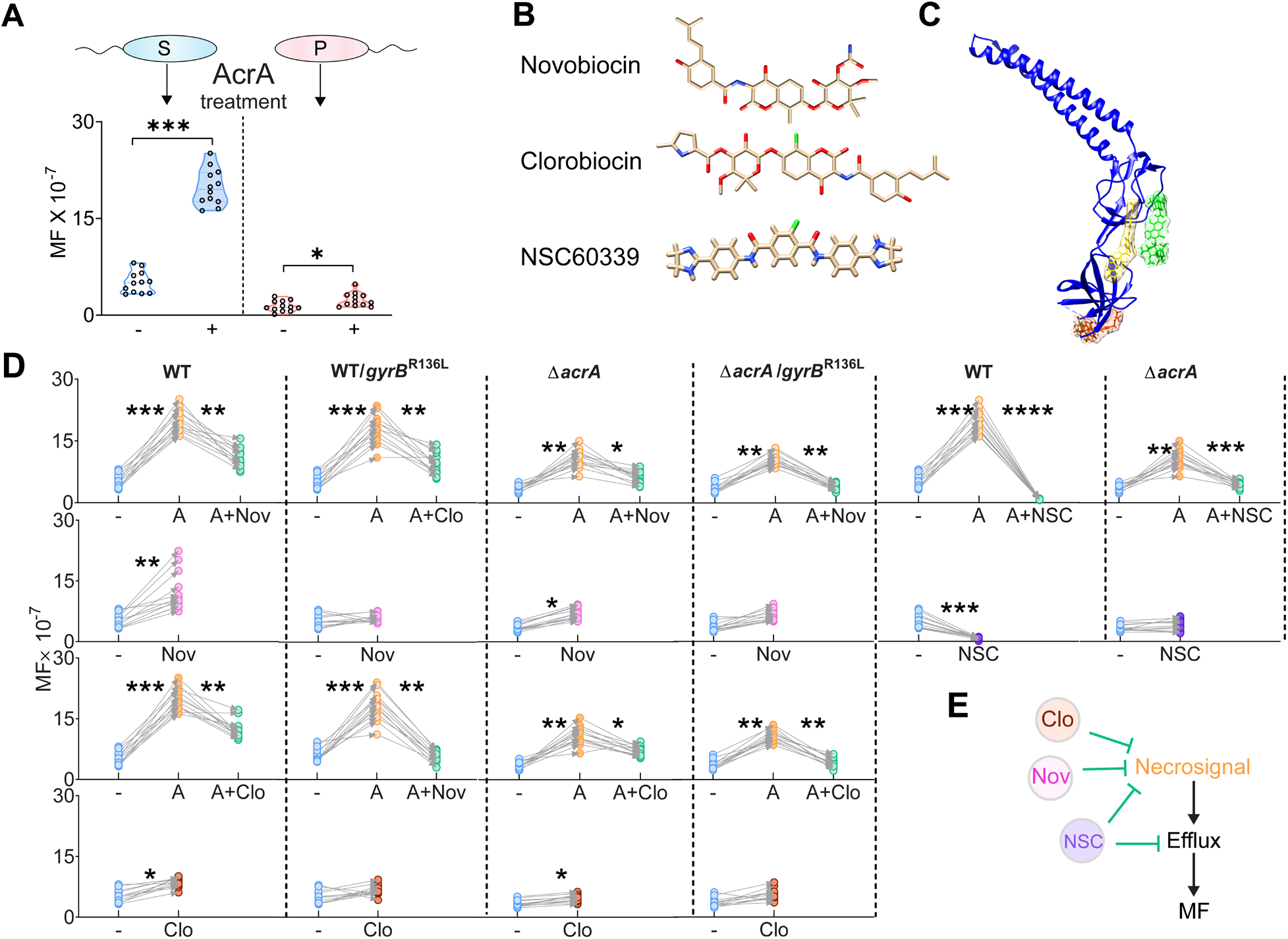
The Mutator Phenotype of Swarms Can be Inhibited by Efflux Pump Inhibitors. (A) Top; experimental design of MF analysis. Bottom; Kan^50^ MF of S (blue) and P (red) treated (+) or untreated (-) with AcrA. [\**p* < 0.05, \*\*\**p* < 0.001, Mann-Whitney test] (B) Potential efflux pump inhibitors. (C) Interaction of inhibitors in B with AcrA (PDB:2F1M). Green: Novobiocin, yellow: Clorobiocin, Red: NSC60339. (D) Rif^50^ MFs of WT and Δ*acrA* strains (S cells) with or without the *gyrB*^R136L^ allele in the presence of necrosignal AcrA and indicated inhibitors. -, no additions; Nov, Novobiocin; Clo, Clorobiocin; NSC, NSC60339; A, purified AcrA. \**p* < 0.05, \*\**p* < 0.01, \*\*\**p* < 0.001, \*\*\*\**p* < 0.0001; Mann-Whitney test. (E) Schematic summarizing the results in D.

In summary, we have shown that drugs targeting both efflux and AcrA reduced the MF of swarms, opening up new possibilities for interfering with evolvability to drug resistance in clinical settings. We note that a wide variety of bacteria, both Gram positive and Gram negative, show necrosignaling (Bhattacharyya et al., 2020). In the case of *E. coli*, the necrosignal AcrA happens to be a component of the efflux pump, making these inhibitors doubly effective. The nature of other necrosignals remains to be investigated.

### A Role for Efflux in Iron Acquisition: Spatial Heterogeneity in Efflux, Redox, and Cell Death

Why do swarms intrinsically exhibit high efflux? Two lines of thought led us to examine whether the primary role of efflux is in obtaining iron from the environment. A variety of studies, including the finding that iron acquisition systems are upregulated, have shown that the bacteria are iron-starved on the surface (Inoue et al., 2007; Lin et al., 2016; McCarter and Silverman, 1989; Partridge and Harshey, 2013a; Wang et al., 2004). Iron-scavenging systems encode high-affinity iron-binding siderophores, which are secreted into the extracellular matrix to bring iron back in (Kramer et al., 2020; Wilson et al., 2016). Efflux pumps are crucial for siderophore secretion (Bleuel et al., 2005; Henderson et al., 2021). If the high efflux activity of swarmers is being employed to obtain iron as the swarm advances into new territory, center and edge zones of a swarm should differ in their amount of siderophores and free iron. Classical staining methods (Schwyn and Neilands, 1987) to monitor siderophores could not be directly used on the swarm plate. We therefore lifted cells from different zones on swarm plates (Figure 6A), to measure siderophores (Schwyn and Neilands, 1987) and iron (Oviedo et al., 2003). The data are plotted as a circular bubble plot to capture the radially symmetric nature of the *E. coli* swarm (Figure 6B-C). Siderophore levels were high at the edge and low at the center, while free iron showed a reverse trend.

**Figure 6.**
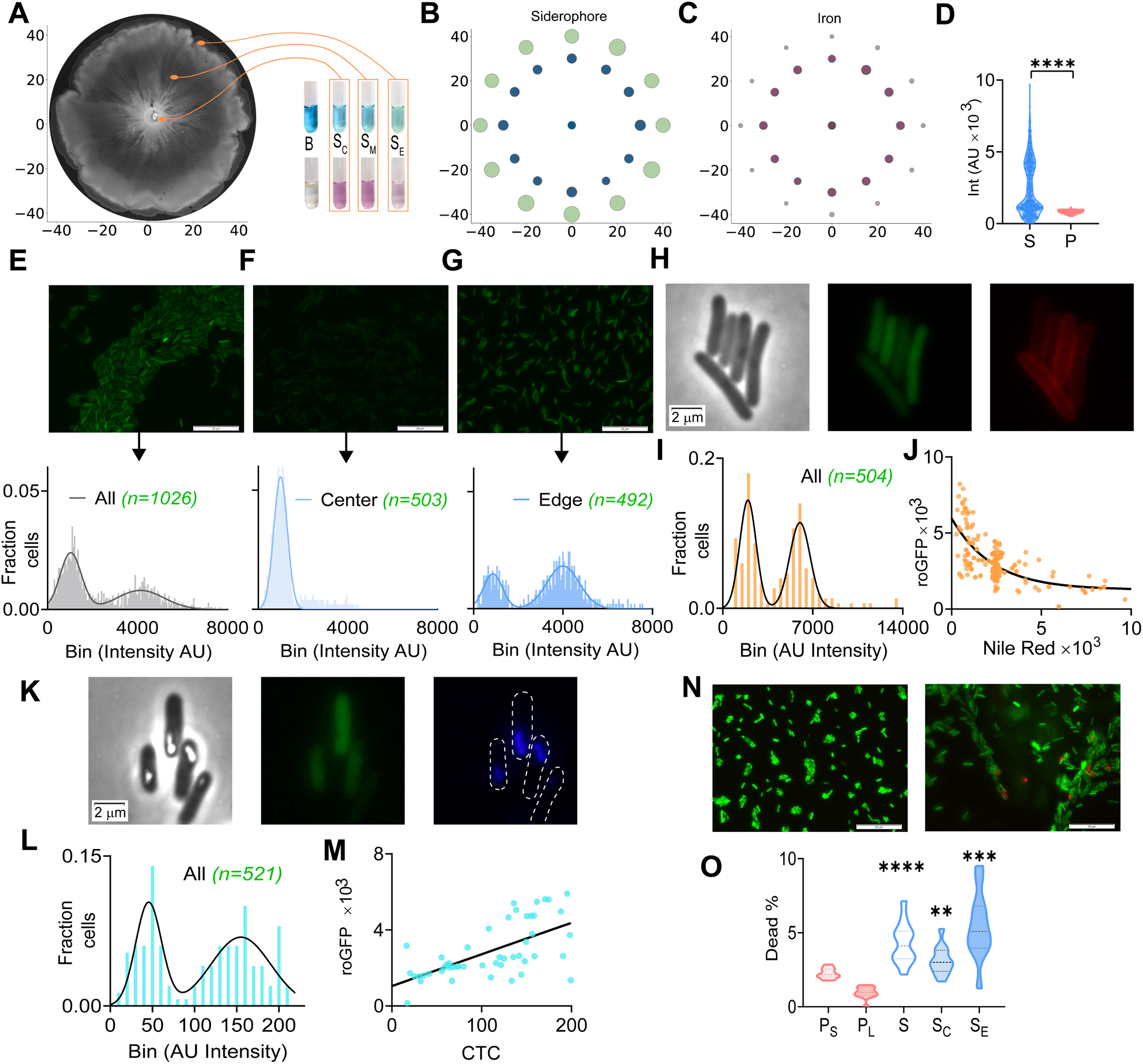
A Role for Efflux in Iron Acquisition: Spatial Heterogeneity in Efflux, Redox, and Cell Death. (A-C) Measurement of iron in swarm cells. (A) Positioning of a swarm plate on a X-Y axis with −45 to 45 mm on each direction from center. Cells from three separate regions were collected, (S_C_: center, S_M_: middle, S_E_: edge, B: blank reagent). (B) Siderophore (blue tubes in A, O-CAS assay) and (C) Iron concentration (magenta tubes in A, ferrozine assay) measurements at the edge and center normalized to the mean of center values. In the circular bubble-plot along the X-Y axis of the plate, each bubble represents one replicate at the indicated locations on the plate. Bubble size and color corresponds to the measured fold-change and color intensity in the assays. (D) Quantification of roGFP intensities in S (n=1026) and P (n=792) cells (see Figure 6E, and Figure S6A for S and P cell data); \*\*\*\**p* < 0.0001; Wilcoxon rank-sum test. (E-G) roGFP fluorescence images (top) and frequency distribution (bottom) of S cells collected from the entire swarm plate. (E) Two populations of fluorescence intensities are observed (AU, arbitrary units) (see Figure S6 C-E for brightfield images). (F, G) as in E except S cells were taken from either the center or the edge. (H-J) roGFP-efflux correlation in S cells. (H) From left to right: bright field, roGFP, Nile red dye. (I) Distribution of Nile red fluorescence. (J) Anti-correlation of Nile red and roGFP (n=202); R^2^= 0.5919. (K-M) roGFP-CTC correlation in S cells. (K) From left to right: bright field, roGFP, CTC (false colored blue). (L) Distribution of CTC fluorescence. (M) Correlation of CTC and roGFP (n=51); R^2^= 0.5012. (N) Live-Dead staining of P (left) and S (right) cells showing superimposed live (SYTO9) and dead (PI) stains (see Figure S6I for brightfield images). (O) Percentage of dead cells. P_S_, P stationary; P_L_, P log phase; S_C_ and S_E_, S cells from center and edge of swarm. [t-tests performed by comparing each S set to P_S_, \*\**p* < 0.01, \*\*\**p* < 0.001, \*\*\*\**p* < 0.0001]

If cells at the advancing edge are engaged in efflux for iron acquisition, would the edge cells experience higher ROS levels and perhaps also cell death as a consequence? Initially, ROS levels were measured in the entire swarm using the redox-sensitive biosensor roGFP, which increases its fluorescence upon oxidation of specific cysteines by ROS moieties inside the cell (Morgan et al., 2011; van der Heijden et al., 2016). S cells were observed to have ~ 6-fold increase in fluorescence intensity compared to P cells (Figure 6D, Figure S6A). The fluorescent signal binned into two distinct sets (Figure 6E) and was noisier (Figure S6B) than that in P cells. The frequency distribution of S cell fluorescence is shown in Figure 6E (graph). We then isolated and measured roGFP intensity of cells separately from the center and edge of the swarm (Figure 6F-G, Figure S6C-E). The center cells showed a single prominent peak of fluorescence indicating a homogeneous population with respect to ROS levels (Figure 6F graph). Edge cells on the other hand were more heterogeneous and displayed two peaks (Figure 6G graph). In addition, edge cells had ~5-fold more mean fluorescence compared to the center cells (Figure S6F).

To test whether the redox heterogeneity is a direct result of efflux activity, roGFP containing S cells were stained with Nile red, which is an indicator dye for efflux (Bohnert et al., 2010) - more Nile red accumulation indicating lower efflux. An anticorrelation between roGFP and Nile red signals was observed (Figure 6H-J, middle and right, and Figure S6G), which followed an exponential decay (Figure 6K). Consistent with the bimodal distribution of ROS profiles, S cells also exhibited a bi-modal distribution in efflux, which indicated the presence of two different efflux cell-types (Figure 5I). A Nile red efflux assay confirmed that edge cells have ~40% higher efflux activity than center cells (Figure S6H). To evaluate whether cellular energetics and respiration are responsible for efflux-driven ROS stress, the redox dye 5-cyano-2,3-ditolyl tetrazolium chloride (CTC) (Rodriguez et al., 1992) was used to stain the roGFP cells. Presence of CTC-formazan is a direct indicator of electron transport chain activity. There was a significant positive correlation between CTC and ROS signals (Figure 6K-M and Figure S6G). Again, the CTC signal revealed two intensity peaks (Figure 6L).

To test the deduction that high ROS causes more death, we treated S cells with a live-dead stain and indeed observed ~4-6-fold higher death within swarms compared to P populations (Figure 6N; Figure S6A). Edge cells specifically had ~3-fold higher death compared to center (Figure 6O).

In summary, high efflux activity at the edge is likely employed for mitigating iron starvation experienced during surface growth, providing an explanation for why S cells engage in more efflux than P cells. In addition to the genetic association between high efflux and an increase in MF, ROS production from efflux itself leads to cellular damage. The data demonstrate that swarm edge and center cells are phenotypically distinct and spatially segregated with respect to efflux and death, with edge cells conducting more efflux and undergoing more death. Cell death is expected to release the necrosignal AcrA, which would further stimulate efflux, and therefore MF (Figure 5A). A high-efflux physiology at the edge would be an added bonus while swarming in relevant habitats where the advancing edge would be expected to be at the forefront of an encounter with antibiotic-producing microbes.

### Phenotype Surfing: an Evolutionary Mechanism for Breeding Mutants that Survive in the Absence of Selection

Thus far, we have seen a striking segregation of various linked phenotypes – increased efflux, high ROS stress, and more cell death – at the edge of the swarm. These phenotypes are also directly correlated with high MF and with evolution of antibiotic resistance mutations. Evolution does not favor high MF because of its deleterious effects in general, especially when the population is dense (Krasovec et al., 2014; Krasovec et al., 2017; Lynch et al., 2016; Sprouffske et al., 2018); in fact, increasing the density of planktonic cultures actually decreases MF (Krasovec et al., 2014). Compensatory mutations that rescue the reduced fitness of such mutants can arise (Bjorkman et al., 2000), but could take a few tens of generations to appear (Sousa et al., 2012), greatly increasing the chances of elimination of the original resistance mutation from the population (Melnyk et al., 2015). So how does the swarm support high MF?

To address this enigma, we first measured the rate of evolution in swarms by applying the concept of ‘evolvability’ E (Pigliucci, 2008), which is the ability of a biological system to produce phenotypic variation that is both heritable and adaptive (Payne and Wagner, 2019). We defined E as a product of mutation frequency (MF) and total size of the population (N). Since N is dependent on the growth rate G of a population, we measured G for both S and P cells (Figure S7A). A 3D plot of these values showed that S cells have an unexpectedly better E than P cells, suggesting that S cells are indeed on a different evolutionary trajectory (Figure 7A). Two observations further support this view. First, we found frequent co-occurrence of two Kan^R^-conferring mutations in S isolates (Figure 1H) in the relatively short selection window of our experiments (24-48 h), an event that is highly unlikely in P cells (Elena and Lenski, 2003), where such mutants were indeed not observed. Second, majority of the Kan^R^ mutants (Figure 1C-D) had a reduced growth rate, hence lowered fitness (Figure 1G).

**Figure 7.**
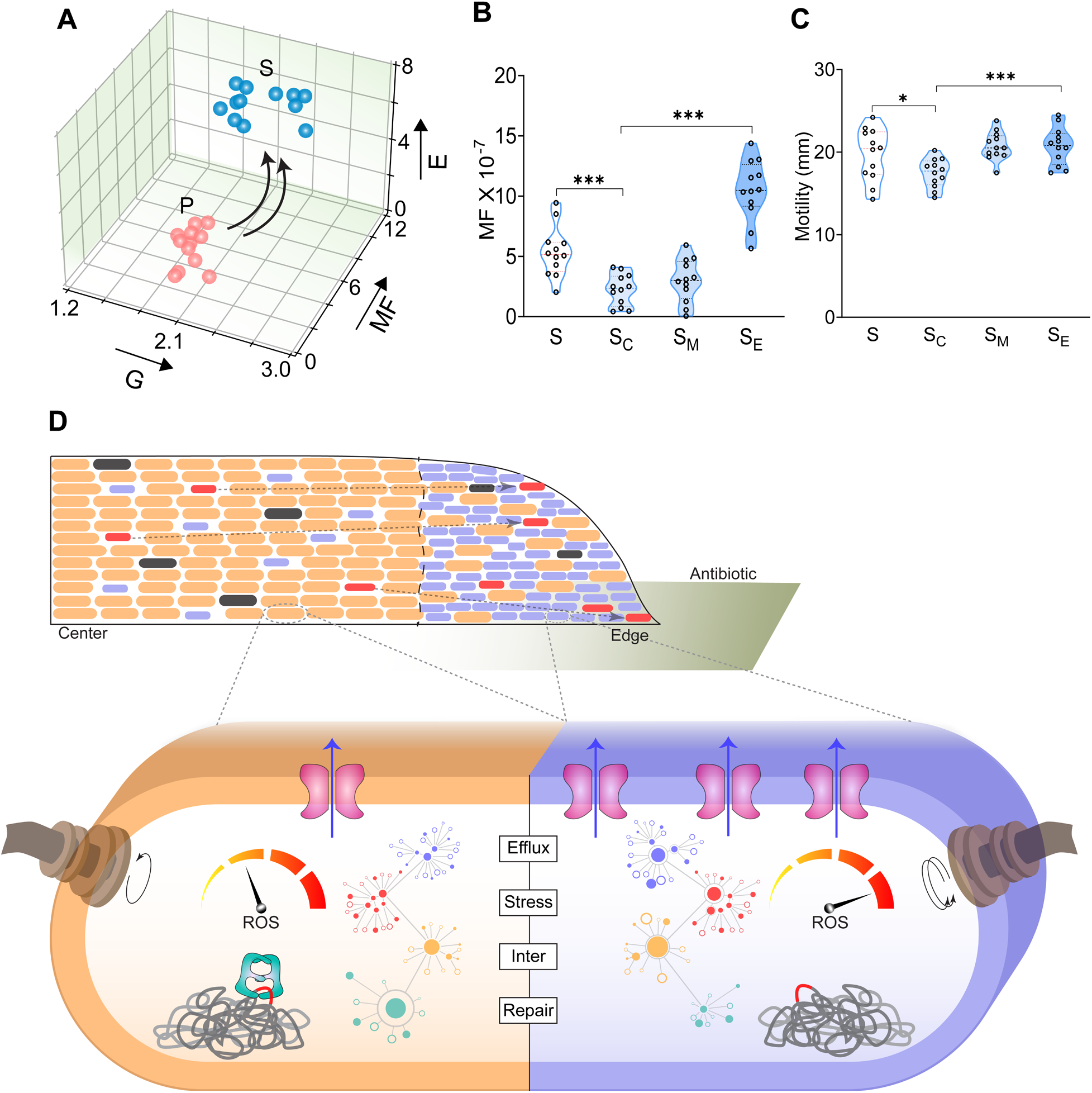
Phenotype Surfing: an Evolutionary Mechanism for Breeding Mutants that Survive in the Absence of Selection. (A) A 3D plot of experimentally derived values of growth rate (G, in generations per hour, x-axis), mutation frequency (MF in 10^-7^/CFU, y-axis), and evolvability (E, Z-axis). Arrows indicate direction of change. (B) Rif^50^ MF of S cells sampled from different regions. S refers to entire population. (C) Swim motility assays with cells collected from swarm plates (see Figure S7G for images). (D) Phenotypic surfing model. Top: a side view of a swarm population with the distribution of two phenotypically distinct cell types (orange and blue) in a spatially segregated edge zone. In this example, the edge has reached an antibiotic zone (olive). Blue cells are more motile and efflux more, producing mutants (red and black) some of which die (black) in absence of a favorable environment. The blue cells and their red progeny ‘surf’ at the edge of the colony either by a faster migration to the edge (dashed arrows) or are favored at the edge to bring in iron. Bottom: Summary of mechanism underlying the phenotypic differences between orange and blue cells. The yellow-red gauge indicates ROS levels, which are high in blue cells. A series of network interactions starting from efflux nodes (blue), through stress (red), and other intermediate nodes (Inter, yellow) decrease DNA repair activity (teal) in blue cells, increasing mutations (red). (D-E) \**p* < 0.05, \*\*\**p* < 0.001; Mann-Whitney test.

Deleterious or neutral mutations that are normally lost by Darwinian forces, have a higher chance of becoming randomly fixed by genetic drift (Lynch et al., 2016), which is high at the edges of an expanding population (Hallatschek and Nelson, 2008). Indeed, it has been shown that favorable mutations can become fixed at the edge via a genetic drift-based mechanism called ‘gene surfing’ (Hallatschek and Nelson, 2008). Using similar principles, we propose a ‘phenotype surfing’ model where the unfavorable phenotype of high MF has an increased chance of survival at the swarm edge. To test this outcome experimentally, we measured the MFs of center and edge cells as before. The data clearly showed that edge cells have ~6-fold higher MF compared to center cells (Figure 7B). In addition, motility within the swarm would affect genetic drift because faster motility would increase the probability of cells reaching the swarm edge (videos S1, S2) where the chances of drift are high (Hallatschek and Nelson, 2008). Indeed, a significantly higher motility was observed in edge cells when compared to center cells (Figure 7C, Figure S7B-G, and ‘Aspect Ratio’ in Supplementary text; see also video S3). The model is also supported by data in Figure 1, where a majority (~70%) of the Kan^R^ mutants isolated from the swarm edge had secondary mutations in *dgcJ*, which specifically enhance swarming motility, implicating their improved motility in transporting Kan^R^ mutations to the front of the expanding swarm. That these mutations occurred within the edge population and were not simply carried there from the center, is attested to by the observation that the mutants were all unique (Figure 1H); had they arisen earlier, one would expect to see many that inherited the exact same Kan^R^-related mutation. Another line of support for the model comes from the observed heterogeneity of fitness costs associated with the Kan^R^ mutations (Figure 1G, S1E, groups 1-4). The data show that mutants with low fitness survived in the swarm despite its dense nature, and were recovered at edge where a favorable selection condition was present – in this case Kanamycin.

Figure 7D summarizes the ‘phenotype surfing’ mechanism derived from the findings reported in this and other sections, and explains how an unfavorable phenotype can increase its survival frequency at the edge. The mechanism is supported by data showing that cells in a swarm are spatially segregated with respect to phenotypes such as MF (Figure 7B) and efflux (Figure S6H) and that motility plays a role in delivering these phenotypes to the edge. Spatial segregation of these phenotypes may be beneficial to the entire colony in two ways. First, the effective population size (Husemann et al., 2016) of a swarm harboring individuals with high genetic relatedness is substantially high. This can support high rates of death as many individuals would still survive. Second, in case of sudden extinction events in the environment that could cause population bottlenecks (Libby and Rainey, 2011), at least some of the spatially segregated individuals would have a better chance of survival. Thus, a perfect storm of events accruing at the edge creates an environment that sustains otherwise unfavorable mutations. The edge phenotype may well be applicable to biofilms if the cells here are engaged in high efflux (Alav et al., 2018). *B. subtilis* biofilms have been demonstrated to have a heterogeneity that includes a special edge function (Larkin et al., 2018).

## CONCLUSIONS

Bacterial swarms provide an excellent platform for studying both the biology and the physics of collective motion and of co-operative behavior. Moving as a dense pack bestows swarmer (S) cells with many survival advantages, but the one we focus on in this study is their temporary capacity to live through antibiotic concentrations that would be lethal if the same bacteria were growing in a planktonic (P) state. At least two swarm features help account for this capacity in *E. coli:* an altered S cell physiology that enables high drug efflux, and a mechanism to further elevate efflux upon encountering antibiotics by necrosignaling. The present study reveals several unexpected facets of the relationship between efflux and DNA repair, of segregation of motile cells with high efflux to the edge of the swarm likely for iron acquisition, of higher cell death and resultant high mutation frequencies (MF) of edge cells. The advancing edge therefore becomes a microcosm for generation and survival of otherwise unfavorable mutations, a novel evolutionary phenomenon we have called ‘phenotype surfing’.

We suggest that the initial condition that sets up the special ecology at the swarm edge is highly motile cells in the population that drive swarm expansion. These cells are also highly metabolically active because motility consumes both PMF and ATP, two energy sources that are also required for efflux. The necessity to acquire iron at the advancing edge therefore favors both phenotypes at the edge because siderophores are secreted via efflux; thus, both phenotypes are favored to expand. Upregulation of efflux brings with it a downregulation of MMR and BER pathways via a network of genetic interactions that appear to respond to the expected ROS stress in the phenotypes segregated at the edge. Dysfunction of DNA repair increases MF. In addition, ROS stress brings with it the risk of cellular damage and collateral death, leading to release of the necrosignal that further increases efflux and hence MF. Mutants that would otherwise have been lost in the interior of the swarm, circulate to the edge and survive there due to decreased competition and increased genetic drift. The positioning of the mutants at the edge provides them an opportunity for range expansion when conditions are ripe, and hence high evolvability to ARMs among other selection forces. Thus, phenotype surfing is a surprising mechanism for evolution in motile bacterial populations, which could in principle also work at the edges of biofilms. In addition to this discovery, our study also provides clues for how to interfere with high evolvability to ARMs.

## ACKNOWLEDGEMENTS

We thank Jeffrey Barrick for critical comments. We also thank Brett Finlay for kindly sending us the roGFP plasmid. This work was supported by Public Health Service Grants GM118085 and AI158295 to R.M.H.

## AUTHOR CONTRIBUTIONS

Conceptualization, S.B. and R.M.H.; Methodology, S.B., and R.M.H; Software, S.B., M.B., A.K.N, and D.M.P; Formal Analysis, S.B., M.B., and A.K.N; Investigation, S.B., D.M.P, and Y.H.; Resources, S.B., M.B, K.H, and A.K.N; Writing – Original Draft and Writing –Review & Editing, S.B. and R.M.H.; Visualization, S.B., M.B, and R.M.H., Supervision and Project Administration, S.B. and R.M.H.; Funding Acquisition, R.M.H.

## DECLARATION OF INTERESTS

The authors declare no competing interests.

## SUPPLEMENTAL INFORMATION

The supplementary information file contains Materials and Methods, Supplementary Text, Supplementary Videos, Figures S1 to S7, Table S1, and References

## Notes

### Competing Interest Statement

The authors have declared no competing interest.

### Summary of Updates

Author order corrected. Text and Figures modified

