## Supplementary Information for "Efflux-linked Accelerated Evolution of Antibiotic Resistance at a Population Edge"

**Brief Contents:**

Materials and Methods

Supplementary Text

Supplementary Videos

Figures S1 to S7

Table S1

References

**Materials and Methods**

**Strains, plasmids, media, and genetics**

Bacterial strains, plasmids, and primers used are listed in supplementary table 1. The strains were propagated on LB broth (10 g/L tryptone, 5 g/L yeast extract, 5 g/L NaCl) or on 1.5% Bacto agar plates. The antibiotic used for marker selection are: Kan (Kanamycin) 25 μg/ml, Amp (Ampicillin) 100 μg/ml, Cam (Chloramphenicol) 30 μg/ml.

For deleting *dgcJ* in *E. coli* MG1655, the one step deletion method was followed by removal of Cam cassette by pCP20 (Datsenko and Wanner, 2000). Other genes were deleted by transferring the Kan^R^ deletion marker from donors in the Keio collection (Baba et al., 2006) using P1 transduction70 (P1vir) (Sternberg and Maurer, 1991). For figure 1J, *dgcJ* was overexpressed from pBAD33. Other genes were overexpressed from ASKA library pCA24N plasmid (Kitagawa et al., 2005) with 0.1 mM IPTG induction.

**Growth and motility assays**

For simplicity, both growth in LB broth and swim motility conditions have been referred to as condition P in the main text. A 0.3% Bacto agar plates were used for swim motility assays by stabbing 4 μL of an overnight bacterial culture at the center of the plates, followed by incubation at 37^o^C. For Figure 7B, the swim plates were incubated for 6h and the rest for 16 h. For Figure 1G, we did not perform growth competition experiments to measure relative fitness (W), as our goal was to evaluate the effect of the resistance mutations even in absence of competition; in addition, multiple passaging would have increased the chances of compensatory mutations. The W was calculated as W = (log(WT)-log(mutant)). OD_600_ at 24h of growth and motility diameter (mm) were used for G and M calculations respectively.

Swarm plates (condition S) were prepared with Eiken agar (Eiken Chem. Co. Japan) for *E. coli* and Bacto agar for others at the following agar concentrations: *E. coli* (0.5 %), *P. aeruginosa* (0.6%), and *B. subtilis* (0.7%). For *E. coli*, 0.5% glucose was included. Swarm plates were dried for 12 h at room temperature before use. Plates were incubated at 30^o^C for *E. coli* and 37°C for others with 4 μL of mid-log phase bacterial culture were inoculated at the center of the plate.

**Isolation and propagation of mutants**

A border-crossing swarm assay setup, as described earlier (Butler et al., 2010) and illustrated in Figure 1B, was used to isolate Kan^R^ mutants. About ~4 μL of mid-log phase bacterial culture (either S or P) were inoculated in the left chamber and allowed to dry for 15 minutes. The plates were then incubated at 30°C for 24-48h. Using sterile loops, mutants from the visible zones of growth on the right chamber were carefully transferred to 5 ml Kan^25^ broth for one-time passaging. A portion of the resultant culture was stored and rest was used for DNA extraction using Wizard genomic DNA purification kit (Promega). The plates were photographed with a Canon Rebel XSI digital camera using the “bucket of light” as a light source (Parkinson, 2007) against a black background such that zones of bacterial colonization appear white and uncolonized agar appears black.

**Genome sequencing**

The genomic DNA of 32 mutants and the parent WT MG1655 were sequenced using the Illumina NextSeq 500 SE75 platform which yielded a total 131.223 million reads. The data was then analyzed to map mutations using the Breseq v0.35.7 platform (Deatherage and Barrick, 2014) against reference genome (GenBank ID U00096.3). The sequencing data were deposited to SRA with BioProject ID PRJNA821886.

**Collection of S and P cells**

S cells from a swarm were collected by flushing the plate with 2 ml LB broth. For S_E_ or edge cells, a 10 mm zone from the edge towards the inside was flushed with LB while the plate was held tilted downward. For S_C_ cells, the central swarm area (10 mm diameter) near the inoculation site (along with the agar at the bottom) was dissected out using a sterile blade and transferred to another sterile petri plate; then, the transferred swarm zone was flushed with LB as before to collect the cells. P_L_ and P_S_ cells (log phase and stationary, respectively) were collected from mid-log phase cells grown till 0.6 and 1.5 OD_600_ respectively.

**Growth rates and Mutation Frequency (MF)**

For growth rates, a single initial colony was inoculated into 24 tubes with 10 ml broth at 37^o^C. 12 tubes were used for P and 12 for S experiments. For P, each overnight culture was subcultured (0.01% inoculum) into fresh media. The doubling time of these cultures was calculated from the log-phase of a growth curve. Simultaneously MFs were calculated by plating 1 ml (adjusted to ~OD_600_=1) culture on each Rif^50^ or Kan^50^ antibiotic hard agar plates and incubated in 37^o^C for 48 hours. All Rif plate work was done under a dark covering over the laminar flow. The resultant colonies were counted and MFs were calculated as resistant colonies/CFU. The results from experimental sets from any day were considered to be usable for interpretation only when the median values of MFs had significant (>95%) CI (Foster, 2006; Kapoor et al., 2019).

For S, each of 12 initial tubes were spotted on 6 swarm plates. Cells from these plates were collected over six time intervals to generate a growth curve. One of these plates was used for collecting cells at the last time point for MF analysis as above.

**DNA damaging agents**

For growth in broth, 200 μl of S and P cells (n=6, adjusted to 0.6 OD_600_) were added to a 96-well plate and treated with 1 mM acidified sodium nitrite in bis-tris buffer pH 5.8 (Nitrite), 20s UV-C ultraviolet exposure (UV), 1 mM hydrogen peroxide (H_2_O_2_), 2 μM bleomycin (Ble), or acidic bis-tris buffer pH 5.8 in 1:100 ratio (Buf). For UV treatment, the microplate was kept open at 2 ft distance from a UV-C source (OSRAM G15T8 OF, 15W Germicidal UV-C Lamp, PURITEC HNS) inside a laminar flow. These plates were kept at 37°C with shaking conditions and OD_600_ was measured after 4 hours. From the same 96-well plate, after the 2-hour treatment, samples were serially diluted and spotted on a rectangular plate.

**Nitrite and peroxide assays**

100 μl of S and P cells (n=12, adjusted to 0.6 OD_600_) were first collected via centrifugation and then washed and resuspended in PBS buffer. Cells were lysed by two freeze-thaw cycles followed by addition of 10 μl of either Measure-IT nitrite reagent (Thermo) or ROS-Glo H_2_O_2_ detection reagent (Promega). The plates were incubated in dark at room temperature for 15 min. Then, 10 μl of signal enhancer or developer reagent was added to nitrite and peroxide samples respectively. For nitrite, the absorbance was then measured with Ex/Em of 365/450 nm; the peroxide samples were measured for their luminescence (RLU). The fold change was calculated setting replicate # 1 from P cells (P_#1_) to one, FC= (S or P)/(P_#1_); the mean of P replicates was not used for normalization in order to observe the variance within P replicates.

**AcrA treatment**

The AcrA protein was purified as reported previously (Bhattacharyya et al., 2020). Each experimental sample was treated with 1 μg of purified AcrA for 2 hours at 30^o^C.

**DNA-efflux correlation**

A previously reported RNA-seq data which compared S to P cells (Bhattacharyya et al., 2020) was re-examined and used to obtain expression of efflux genes (Ecocyc, (Keseler et al., 2011)) and DNA repair genes (REPAIRtoire (Milanowska et al., 2011)).

**Microscopy**

All cells were visualized using a light microscope with fluorescent channels (BX53F; Olympus, Tokyo, Japan) and recorded using cellSens software (v1.6).

Live-dead staining was performed with LIVE/DEAD™ BacLight™ Bacterial Viability Kit (Thermo). Cells were treated with 0.3% red and green dye mixture, and incubated for 15 min in dark at room temperature. All cells are stain with the green dye SYTO9, but only membrane damaged dead cells stain with the red dye propidium iodide (PI). 100 cells from 12 replicates of each sample were counted.

To measure intracellular ROS, the fluorescence intensities of S and P cells, expressing roGFP2 protein from pfpv25.1 plasmid (Staudacher et al., 2018; van der Heijden et al., 2015), were recorded using a Chroma 405 filter with ET405/20x excitation, T425LPXR dichroic, and ET510/40m emission. The signals intensities from a GFP filter (Ex, 460–480 nm; Em 495–540 nm), which showed very little variability, was used for normalization of the 405 signals. To measure efflux, 100 μl of S and P cells harboring pfpv25.1/roGFP2 plasmid were treated with 10 mM Nile Red (VWR) and shaken at 200 rpm for 30 min at 37 °C (Bhattacharyya et al., 2020); under the microscope, the Nile Red intensity was recorded using a filter with EX 545/30x excitation, T570lp dichroic, and EM620/60m emission. To measure active respiration using CTC (Rodriguez et al., 1992), 100 μl of S and P cells harboring pfpv25.1/roGFP2 plasmid were treated with 2mM CTC (VWR) and shaken at 200 rpm for 10 min at 37 °C. Under the microscope, the CTC intensity under the microscope was recorded using a filter with EX 545/30x excitation, T570lp dichroic, and EM620/60m emission. Microscopic images were processed using the MicrobeJ tool (Ducret et al., 2016) in ImageJ package (Schneider et al., 2012) to measure fluorescent intensities of each cell. For segmentation mask (Panigrahi et al., 2021), the following shape descriptors were kept: area [p^2^] = (350-2000), length[p] = (0.5-max), width[p] = (0.1-max), and angularity = 0-1. Using a threshold of 500 units, the cells were segmented and normalized intensities were measured in arbitrary units (AU) and frequencies were counted in 50 AU bins (0-8000). Mean Fluorescence Intensity (MFI) per area in a cell was measured as = (fluorescence in AU)/ (total pixels in a cell). MFI per cell was measured as: (MFI x cell number)/ (mean area of a cell). A non-linear regression was used for curve fitting in Figure 6E-H, K with sum of two gaussians with constrains (0 ≥ mean1 ≥ 30, mean2 ≥ 30). The kurtosis for the aspect ratio of cells were calculated as k= (μ_4_/σ^4^) where μ_4_ is the fourth central moment and σ is the standard deviation.

**Siderophore and iron assays**

Siderophores were detected using methods described previously (Perez-Miranda et al., 2007; Schwyn and Neilands, 1987). Briefly, using the broad end of a 200 μl microtip, zones of swarm containing both S cells and the agar beneath were sampled from different positions on the plate. The collected material was then resuspended in 2 ml CAS reagent (0.5% Chrome azurol S dye, 0.15% and hexadecyltrimetryl ammonium bromide, 0.7% PIPES; pH adjusted to 6.5) and incubated in dark for 4 hours. The A_430_ of samples were measured against the blank. For estimating iron, samples were collected as mentioned in the siderophore section and then resuspended in ferrozine detection reagent (0.2% ferrozine dye in 50mM acetate buffer, pH adjusted to 6 using 10 mM HCl) ((Oviedo et al., 2003). The samples were incubated in dark for 4h. The A_560_ of samples were measured against the blank.

**Treatment with inhibitors**

1 ml of S cells (n=6, adjusted to 0.6 OD_600_) were first collected via centrifugation and then washed and resuspended in LB. Then the cells were treated with 1 μg/m Nov (novobiocin, VWR), 1 μg/ml Clo (clorobiocin, AdooQ), 1 μg/ml NSC (NSC60339, MedChemExpress), and purified AcrA (1 μg/ml) for 2 h followed by MF analysis.

**Nile Red efflux Assay**

The assay was performed as reported previously (Bhattacharyya et al., 2020) with modifications. Briefly, cells were incubated with 10 mM CCCP for 30 min at RT, then 10 mM Nile Red was added and the culture was incubated at 37 °C for 30 min with shaking (200 rpm). These cells were kept in room temperature for 15 min without shaking and harvested via centrifugation. The resultant pellet was resuspended in PPB to obtain OD_600_ of ~1.0 and diluted 10× which then was transferred to a quartz cuvette to measure fluorescence (excitation at 552 and emission at 636 nm) using Spectramax M3 spectrometer every 10 s for 100 s. The Nile Red efflux was initiated by addition of 100 μl of 1 M glucose; fluorescence was measured every 10 s for 200 s. The obtained A_636_ was normalized with OD_600_.

**Structural analysis**

All the structures were visualized using Chimera (Pettersen et al., 2004). In case of EF-G [PDB:4V7D (Brilot et al., 2013)], the Kan moiety was mapped to the structure guided by PDB:2ESI (Huang et al., 2016). The inhibitors (novobiocin: green, Pubchem CID 54675769; clorobiocin: yellow, Pubchem CID 54706138; NSC60339: red, Pubchem CID 65558) were docked using Chimera guided by previous studies (Abdali et al., 2017; Darzynkiewicz et al., 2019).

**Statistics and visualization**

The statistical analyses were performed using Prizm v8 (Graphpad), R studio, Python (ggplot and numpy packages) using Jupyter, and Microsoft Excel. The data was visualized in Prizm, Gnuplot, and python.

**Supplementary Text**

**Network Analysis**

The entire genetic network of *E. coli* was obtained from Ecocyc (Keseler et al., 2011). Then, starting from the DNA repair genes (*mutS, mutM,* and *ung*) and efflux genes, a network of interest was extracted using a bottom-up approach, which included all second-level unique interactions (Figure 3D, top). Peripheral nodes which had only one connection, were filtered out (Figure 3D, bottom). Next, all the paths that connect the selected efflux and DNA repair genes were identified. Two consecutive nodes in those paths were considered as a gene pair. For first level of validation, the COLOMBOS database (Moretto et al., 2016) was used to obtain curated and normalized *E. coli* gene expression data from 4000 experimental sets. The fold change values for each gene in the pair were obtained from entire expression datasets (Figure 3B; plotted as ±10-fold X-Y coordinates which covered all but 0.0025% of the values). The values where one gene of a pair showed significant fold change (± 1.5-fold), the expression value of the other gene in that same pair was used to calculate the Pearson correlation coefficients (CC) of expression. For a statistical comparison (±0.3 cutoff), 100 random gene pairs, generated with a conditional probability randomization (Quicksort), were used. For visual comparison, the CC from the first random pair has been shown in the step plot (Figure 3C, top) and rest are shown in Figure S3A. Only the significant pairs (see Figure S3B for rejected paths) were further filtered using our swarm specific RNA seq data (further supported by genetic experiments, see main text). The final regulatory network was mapped and visualized using Cytoscape v3.8.0. The PERL and Shell scripts are provided separately in supplementary file 1.

**ARM analysis**

The ARM data were collected from CARD server (Alcock et al., 2020). The data specific to *E. coli* were further used to classify the ARMs into categories based on highest level GO processes. The number of ARMs belonging to each class was counted for each genome. If any genome contained at least one efflux ARM, it was categorized under ‘efflux ARM genome’ and so on for each category. A Spearman correlation coefficient matrix was generated comparing all categories in pairs. The PERL script has been provided separately in supplementary file 1. For statistical comparison of correlation coefficients, COCOR package (Diedenhofen and Musch, 2015) was used to generate z-scores for each coefficient pair (Hittner et al., 2003) taking zero difference between them as null hypothesis.

**Aspect Ratio**

Two recent studies – both experimental and theoretical – observed that within a swarm colony, highest speeds are obtained not by largest or smallest cells but cells of intermediate size (Bera et al., 2021; Ilkanaiv et al., 2017). To test if this holds true in our data, we measured the cell morphologies of S cells. The edge and center populations were different in their morphology, with center cells being longer and having a higher aspect ratio (ratio of width and length) than edge cells (Figure S7B-F). Interestingly, the distribution of aspect ratio of center cells deviated from a Gaussian distribution showing significantly high kurtosis values (Figure S7E; kurtosis defines how heavily the tails of a distribution differ from normal). This would mean that the aspect ratio of our edge cells is presumably ideal (~3:1), and these cells would move faster than center cells with a higher aspect ratio (3.8) (Figure S7B-F).

**Supplementary Videos**

*E. coli* swarm plates were placed on the stage of a Nikon Eclipse Ni compound light microscope and visualized with a 40X dry (Plan Fluor 0.75 NA, Ph2) objective. Videos were recorded on a Nikon DS-Ri2 camera with 50 ms exposure, and the black and white colors inverted for better contrast first using Adobe Premiere Pro software and then encoded with H.265 (HEVC) codec. On average, the swarm cells are ~4 μM long in all the videos.

**Video S1. Range expansion at a swarm edge**

10 sec clip of the moving edge of a swarm colony in real-time. The monolayer of cells at the very edge is being pushed forward by cells immediately behind.

**Video S2. The swirling middle zone in a swarm**

Cell density is higher here compared to the edge, creating swirling vortices as described previously (Harshey, 2003; Partridge et al., 2018).

**Video S3. The dense center of a swarm**

The center is at the initial site of inoculation. Cells are densely stacked on top of each other and very little motion is evident.

**Supplementary Figures**

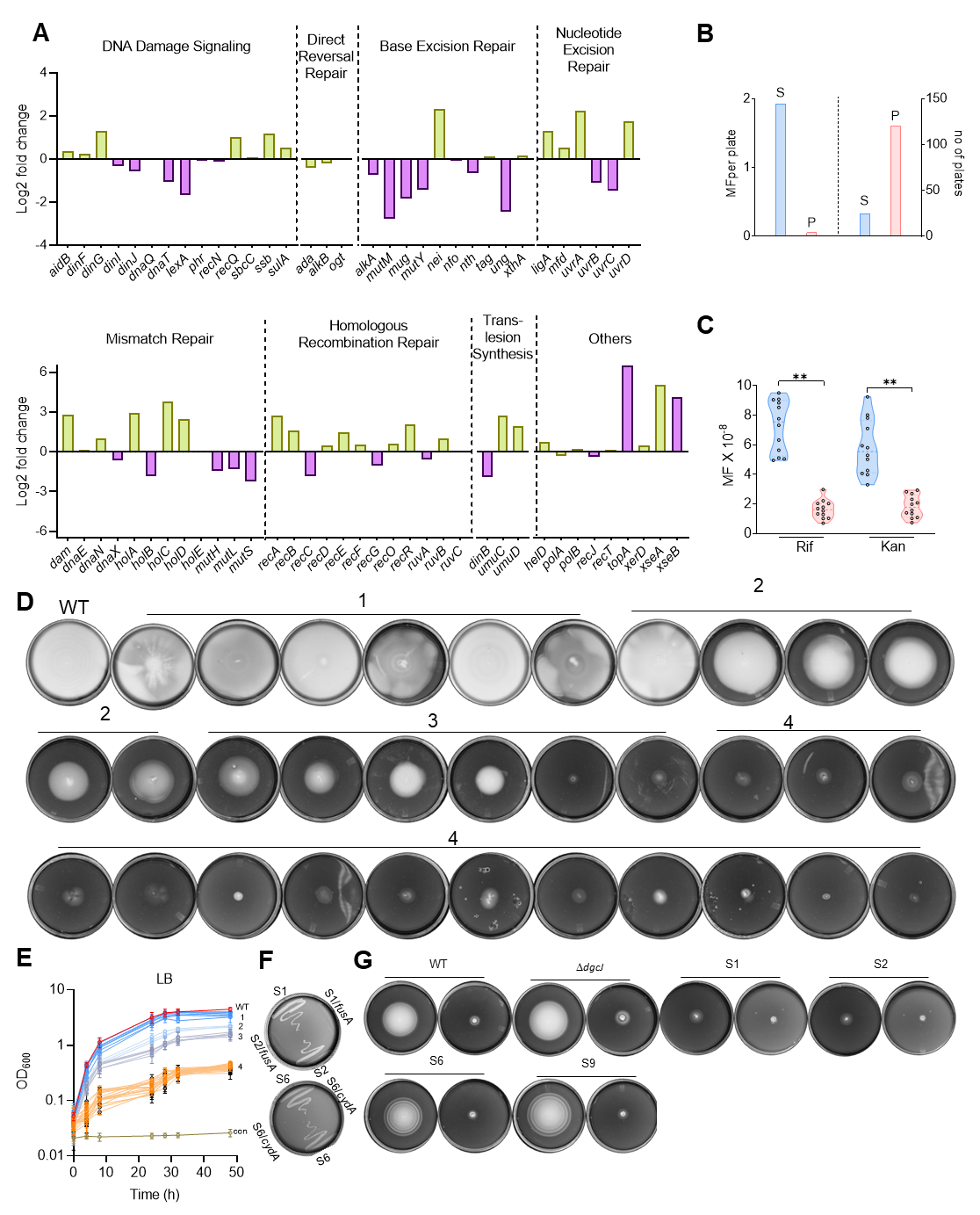

**Figure S1.** **Swarms Downregulate DNA Repair and Show High Evolvability to Antibiotic Resistance**

(A) Mean fold changes in gene expression of different classes of DNA repair genes (Bhattacharyya et al., 2020). Figure 1A only highlights those downregulated more than 1.5 log2-fold.

(B) Quantitation of data in Figure 1C-D. Left, frequency of mutants isolated per plate. Right, total number of plates. S mutants were found in 12/25 plates, and P mutants in 5/121 plates.

(C) Mutation frequencies (MF) of S (blue) and P (red) cells of *P. aeruginosa*. See Figure 1E-F for details.

(D) Representative motility on soft agar 'swim' plates of WT and S-P mutant groups 1-4 which were used to generate heat map shown in M of Figure 1G.

(E) Growth curve of WT and P mutant groups in LB which was used to generate heat map shown in G of Figure 1G; con: LB control for contamination.

(F) Growth of *fusA* and *cydA* Kan^R^ mutants S1, S2 and S6 (Figure 1G-H) on Kan^50^ plates to demonstrate that overexpression of *fusA* and *cydA* in these strains abolishes Kan^R^.

(G) Representative swim motility of *dgcJ* Kan^R^ mutants S1, S2, S6 and S9 (Figure 1G-H) along with WT and Δ*dgcJ* controls. All strains were complemented with a DgcJ-encoding plasmid under pBAD control. In each pair of plates, arabinose was added only to the plate on the right.

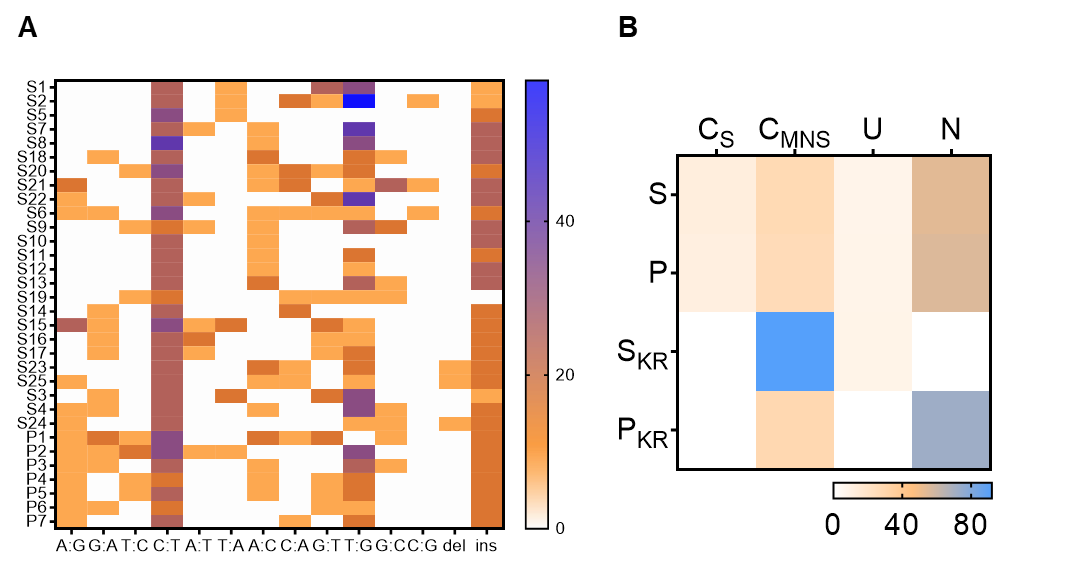

**Figure S2. High Mutation Frequency in Swarms is Caused by a Deficiency in Specific DNA Repair Pathways**

(A) Mutation spectrum of all S and P isolates. The nature of nucleotide changes is indicated on the x-axis.

(B) Heatmaps showing percentages of different classes of mutations found in all S and P mutant isolates based on genic location. Coding region in whole genome with sense (C_S_) or mis-/non-sense (C_MNS_) mutations, upstream of ORF (U), and non-coding (N). S: S whole genome, P: P whole genome. KR: Kan resistant mutations only.

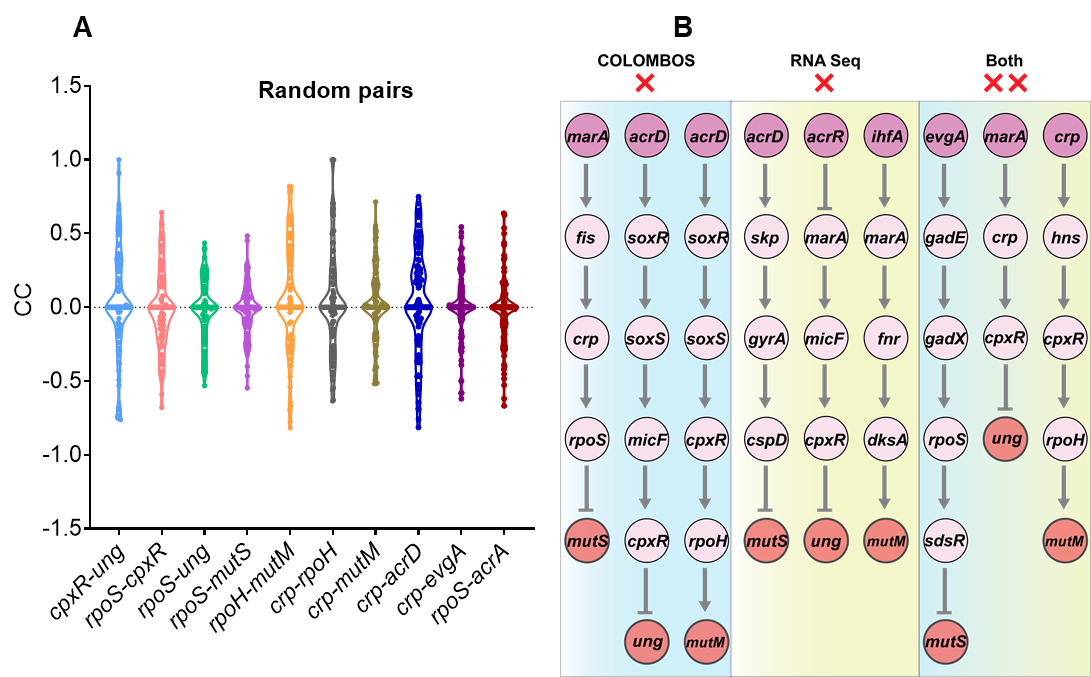

**Figure S3. A Global Regulatory Connection Between Efflux and DNA Repair**

(A) The correlation coefficient (CC) values of random gene pairs generated for statistical analysis for each gene-pair of interest indicated at the bottom.

(B) From the network shown in Figure 3D (Top), the paths with non-significant CC in the COLOMBOS dataset were rejected (x) first. Then, RNA seq data was also used to validate these paths and were rejected if non-significant fold changes were observed. Some examples of paths which were rejected by both approaches are shown on the right.

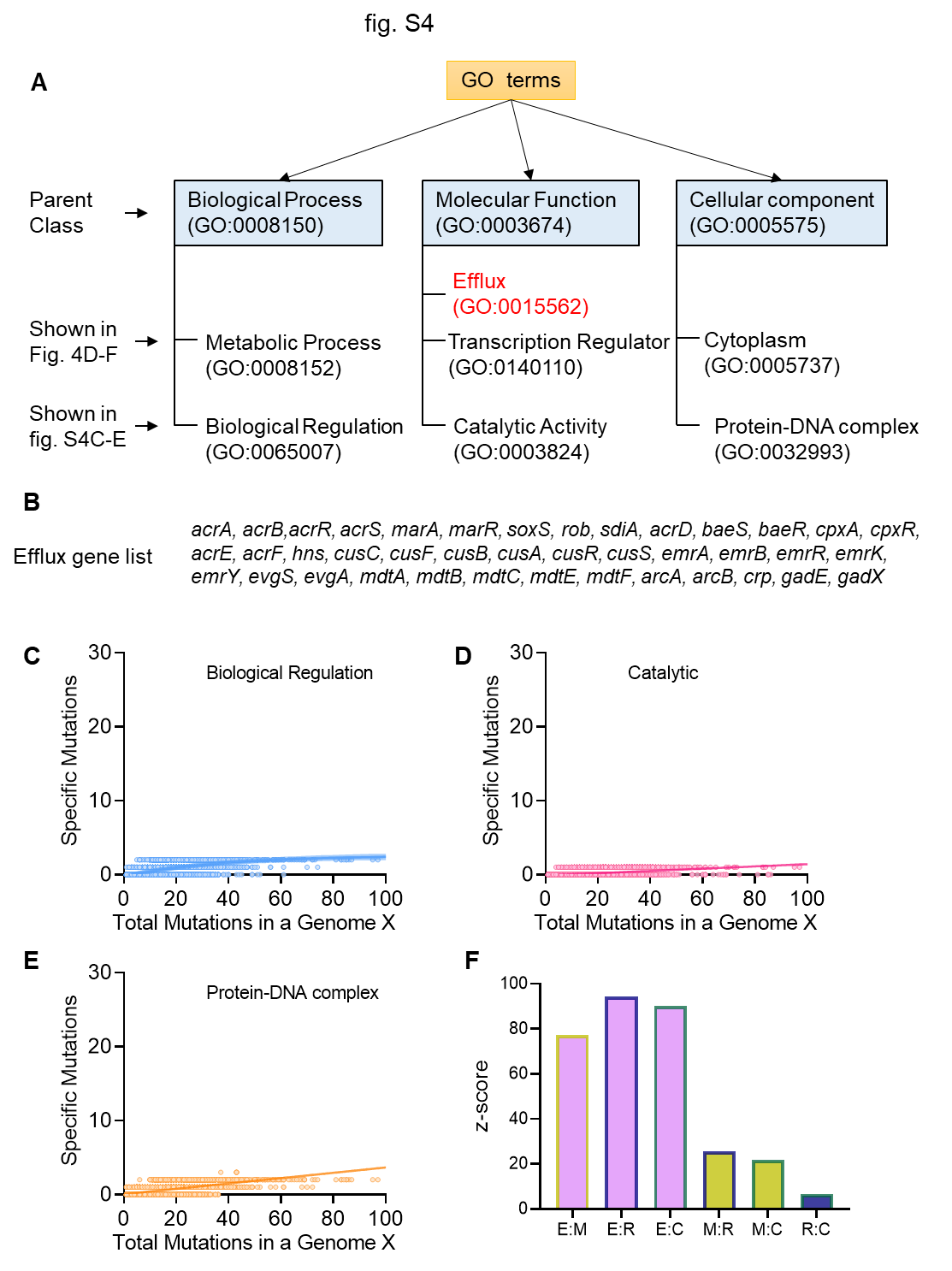

**Figure S4. The Efflux-DNA Repair Network is Clinically Relevant to the Rise in Antibiotic Resistance**

(A) The GO terms used for the analysis reported in Figure 4A-G where data for only efflux (GO:0015562), metabolism (GO:0008125), regulation (GO:0140110), and Cytoplasm (GO:0005737) are shown.

(B) Efflux gene list obtained from Ecocyc.

(C-E) Scatter plots of number of mutations in a specific category in a given genome X vs number total of mutations in of genes in that genome.

(F) Significance of the differences in specific pairs of correlation coefficients (CC) in terms of z-scores. E: efflux-total; M: metabolism-total; R: regulatory-total; and C: cytosolic-total. For example, a higher z-score for the E and M correlation pair (E:M) than the M and R pair (M:R) shows that the CC value of E is significantly high than that of M. The z-score was calculated using COCOR package (see ‘Database Analysis’ in Supplementary Text).

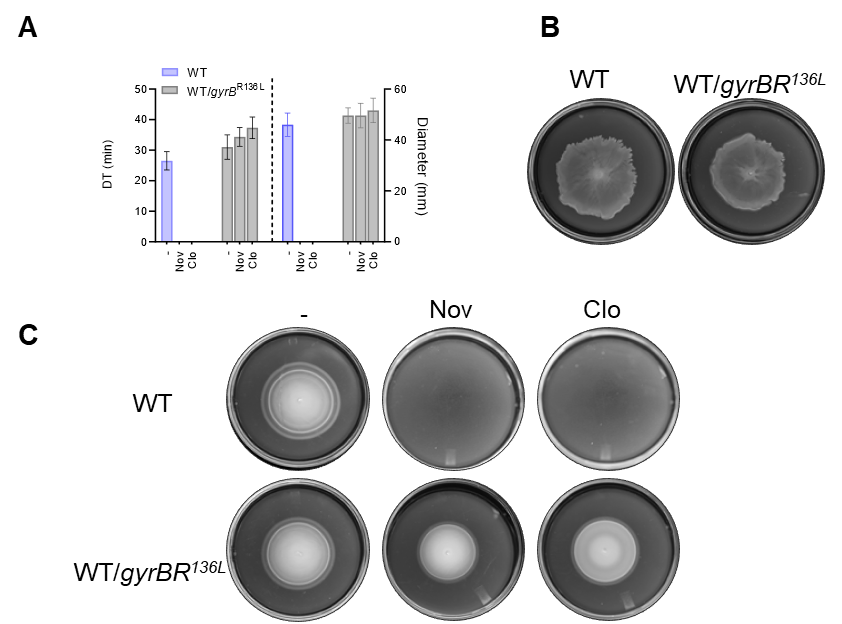

**Figure S5. The Mutator Phenotype of Swarms Can be Inhibited by Efflux Pump Inhibitors**

(A) Growth (DT, doubling time, left) and motility (right) of WT *E. coli* and WT expressing *gyrB*^R136L^ allele, in absence or presence of 1 μg/ml of indicated inhibitors.

(B) Swarm plates showing that overexpression of the mutant gyrase had no significant effect on the swarming ability of WT cells.

(C) Swim motility plates of strains with or without the presence of compounds as indicated on top.

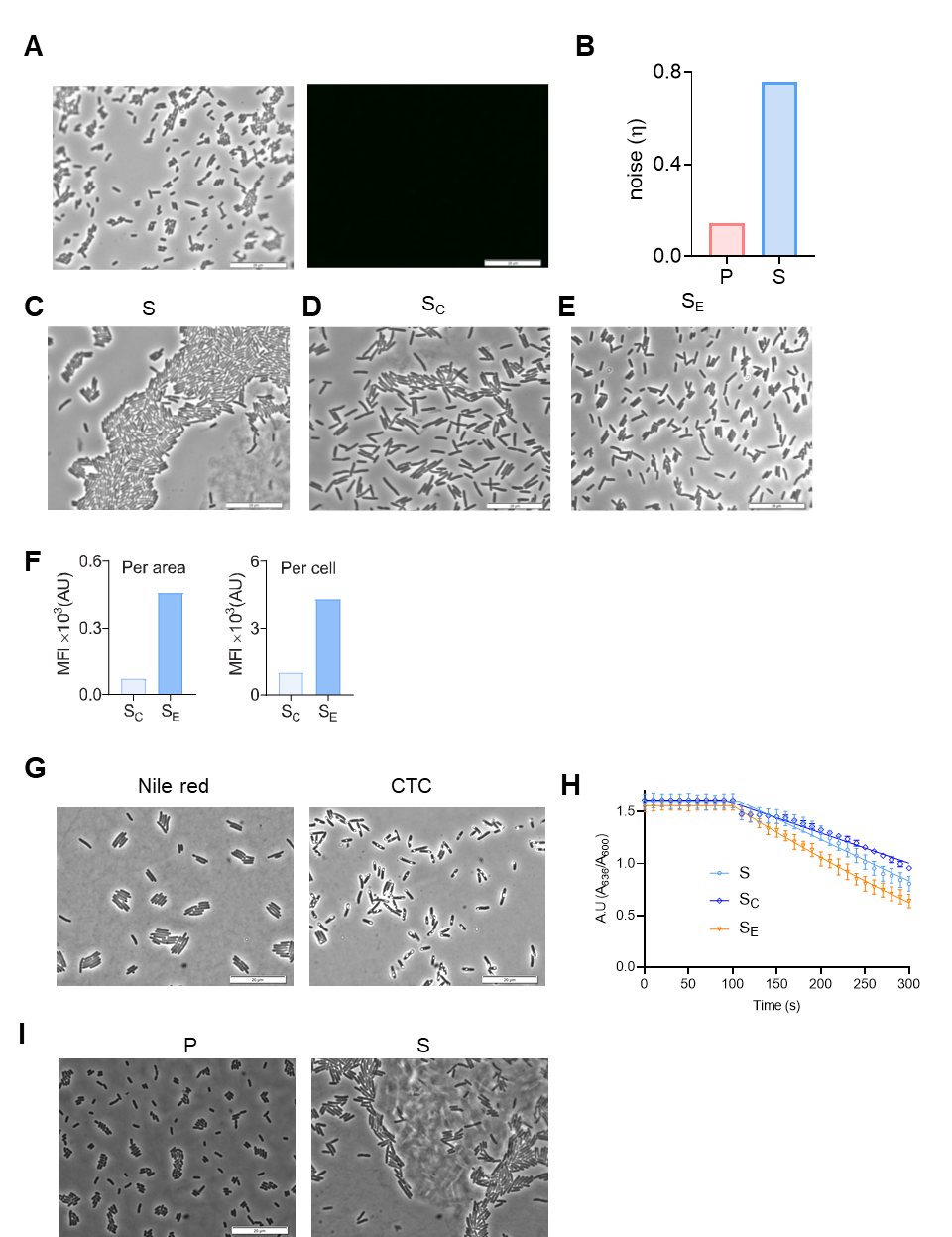

**Figure S6.** **A Role for Efflux in Iron Acquisition: Spatial Heterogeneity in Efflux, Redox, and Cell Death**

(A) Brightfield images (left) and roGFP fluorescence (right) of P cells.

(B) Noise calculation for Figure 6D where noise is defined as $\eta=(SD/M)$ where SD is standard deviation of the samples, and M the population mean. An increase in $\eta$ indicates more heterogeneity within a population

(C-E) Brightfield images for Figure 6E-G.

(F) Mean fluorescence intensity (MFI) per cell (left) or per unit cell area (right) of all cells in each sample category.

(G) Brightfield images for the entire field, only a subset of which were shown in Figure 6H and K.

(H) Efflux assay using Nile red indicator dye (n = 3). Glucose was added at 100 s to initiate efflux. Nonlinear regression analysis of indicated samples (see key). The rate of efflux was measured using the following formula: [(Initial AU at 100s) - (Final AU at 300 s) / (200 × (Initial AU at 100s)). The obtained rates were: S = 2.48 ×10^-3^, SC = 2.01 ×10^-3^, SE = 2.93 ×10^-3^ in AU s^-1^ units.

(I) Brightfield images for Figure 6N-O.

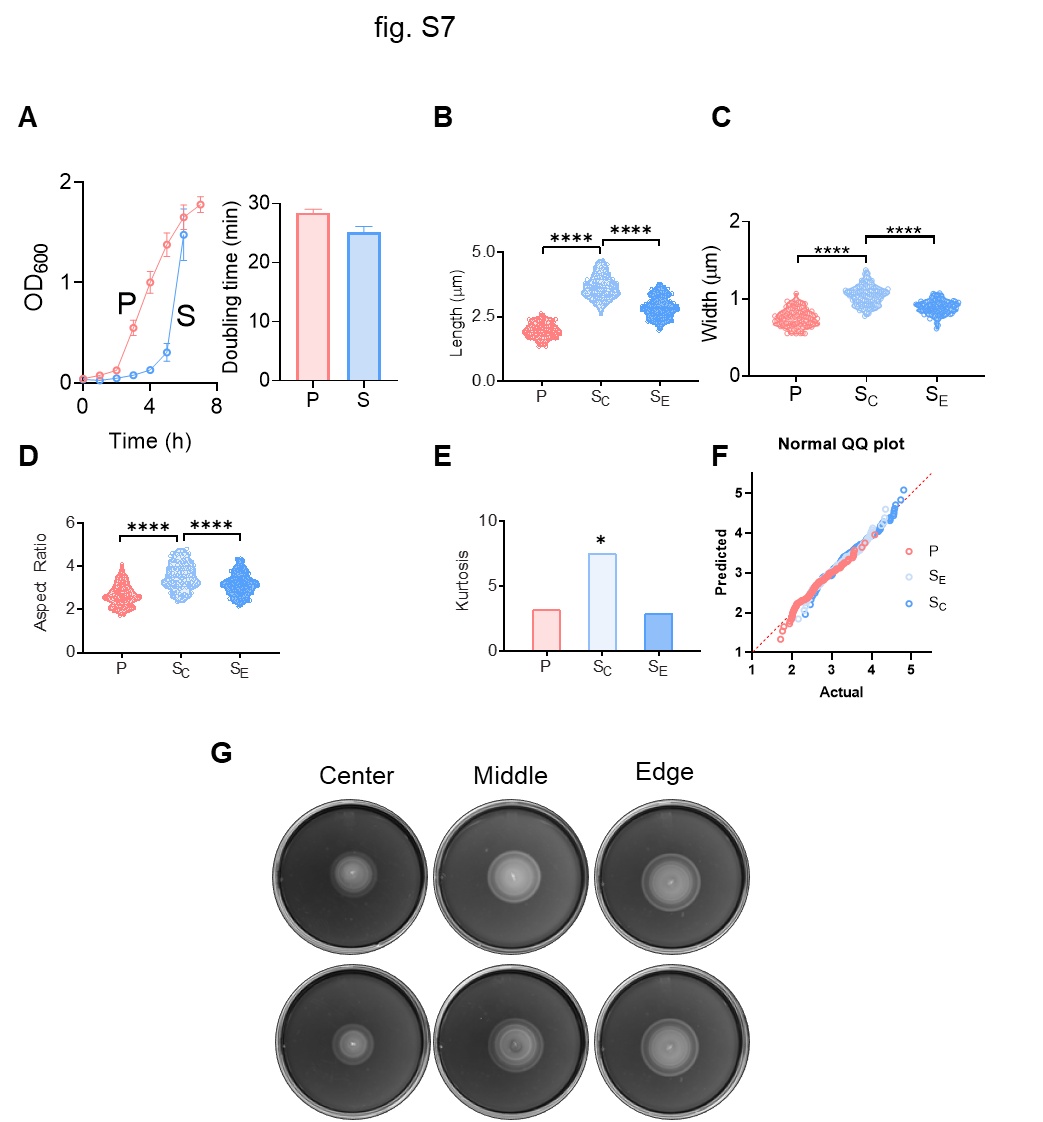

**Figure S7. Phenotype Surfing: an Evolutionary Mechanism for Breeding Mutants that Survive in the Absence of Selection**

(A) Growth curves of S and P cells (left) and calculated doubling time (right).

(B-E) Comparison of cell morphologies of P and S cells, the latter taken from either the center (c) or edge (e) (n=100). Cell-length (B), width, (C), their length-width aspect ratio (D), and D'Agostino & Pearson normalcy test on the distributions shown in E (E). Kurtosis (k) is a measure of the deviation from Gaussian distribution of cells [**p* < 0.05].

(F) A quantile-quantile (QQ) plot of aspect ratio from Figure S7E; as another measure of the normalcy of a data, the actual residual is plotted against predicted residual from a Gaussian distribution. Among the three, only S_C_ shows over-dispersed nature on both tails and did not pass the normalcy test.

(G) Swim motility plates of cells from different positions of swarm. The two rows are two replicates.

**Table S1. Strains, Plasmids, and Primers**

| **Strain/Plasmid** | **Genotype** | **Source and/or reference** |
| --- | --- | --- |
| **Strains** | | |
| ***Escherichia coli*** | | |
| MG1655 | Wild type; F^-^ λ^-^ *rph*-1 | (Guyer et al., 1981; Partridge et al., 2015) |
| Δ*dgcJ* | MG1655 Δ*dgcJ::*Kan^R^ | This study |
| Δ*marA* | MG1655 Δ*marA::*Kan^R^ |  |
| Δ*evgA* | MG1655 Δ*evgA::*Kan^R^ |  |
| Δ*marA*Δ*evgA* | Δ*marA* (kan removed), Δ*evgA::*Kan^R^ |  |
| Δ*acrD* | MG1655 Δ*tolC::*Kan^R^ |  |
| ΔacrA | MG1655 Δ*tolC, Kan^R^* removed |  |
| ***Bacillus subtilis*** |  |  |
| 3610 | Motile wild type | (Kearns and Losick, 2003) |
| ***Pseudomonas aeruginosa*** | | |
| PAO1 | Wild type | Lab collection |
| **Plasmids** | | |
| pCA24N | From ASKA collection, Cam^R^ | ASKA (Kitagawa et al., 2005) |
| p*fusA* | pCA24N carrying *fusA* |  |
| p*cydA* | pCA24N carrying cydA |  |
| p*dinB* | pCA24N carrying *dinB* |  |
| p*holA* | pCA24N carrying *holA* |  |
| p*lexA* | pCA24N carrying *lexA* |  |
| p*mug* | pCA24N carrying *mug* |  |
| p*mutS* | pCA24N carrying *mutS* |  |
| p*mutM* | pCA24N carrying *mutM* |  |
| p*ung* | pCA24N carrying *ung* |  |
| p*recC* | pCA24N carrying *recC* |  |
| p*katE* | pCA24N carrying *katE* |  |
| p*sodB* | pCA24N carrying *sodB* |  |
| p*katG* | pCA24N carrying *katG* |  |
| p*marA* | pCA24N carrying *marA* |  |
| p*evgA* | pCA24N carrying *evgA* |  |
| p*feoB* | pCA24N carrying *feoB* |  |
| pBAD24 | Cloning vector, P_BAD_, Cam^R^ | (Guzman et al., 1995) |
| pBAD33 | Cloning vector, P_BAD_, Amp^R^ |  |
| pBAD*dgcJ* | pBAD33 carrying *dgcJ* gene | This study |
| pfpv25.1 | Cloning vector, Amp^R^ | (van der Heijden et al., 2015) |
| pfpv25.1/roGFP2 | pfpv25.1 carrying roGFP2 |  |
| pTrc99a | Cloning vector, P_Trc_ and Amp^R^, | (Amann et al., 1983) |
| *gyrB*^R136L^ | pTrc99a carrying mutant *gyrB*^R136L^ | This study |
| pCP20 | Rep^ts^, Amp^R^, Cam^R^ | (Cherepanov and Wackernagel, 1995) |
| p*acrA* | pTrc99a carrying WT acrA with N-terminal His-tag and C-terminal FLAG-tag | (Bhattacharyya et al., 2020) |
| **Primers** | | |
| **Name** | **Sequence (5’🡪3’)** | |
| acrA KO fp | TACCATAGCACGACGATAATAT | |
| acrA KO rp | ATCACCCACGCAAAAATCGGG | |
| marA KO fp | AACAAAAAACCTGACGGCGG | |
| marA KO rp | TGCGCGGAAAAGAGAATAAG | |
| evgA KO fp | TATTGAGAAAATGAGATGAC | |
| evgA KO rp | CGAAACTTATGGTCGACCAA | |
| acrD KO Fp | GTCAGTTAATGTAATGCCTCCTAC | |
| acrD KO Rp | TACAAACAGCAAGAACCCGC | |
| acrB KO Fp | GAAAGTGCGTCCTGGTGTCCAG | |
| acrB KO Rp | TATGAGATCCTGAGTTGGTGG | |
| dgcJ Datsenko Fp | GGTTTTCTCGTTTCACTAACCGAAGGAGTGCCATTTATCTGTGTAGGCTGG  AGCTGCTTC | |
| dgcJ Datsenko Rp | CACGCTCCTGAGATTACAAGCAAACAACCACAGAAGGTTACATATGAATAT  CCTCCTTA | |
| dgcJ pBAD Fp | TAAGCAGAGCTCAGGAGGAATTCACCATGAAATTGCACCATAGAAT | |
| dgcJ pBAD Rp | TGCTTATCTAGATTATCATGAACGGCTGTTTTTGT | |
| GyrB Fp | CGAGAAGGATCCATGTCGAATTCT | |
| GyrB Rp | GCTCGCAAGCTTAGCGCCATTAAA | |
| R136L Fp | CCCTGTCGCAAAAACTGGAGCTGGTTATCCAGCTGGAGGGTAA | |
| R136L Rp | CCGTGTTCGTAGATCTGACGGTGAATTTTACCCTCCAGCTGGATA | |
